# An interaction map of transcription factors controlling gynoecium development in Arabidopsis

**DOI:** 10.1101/500736

**Authors:** Humberto Herrera-Ubaldo, Sergio E. Campos, Valentín Luna-García, Víctor M. Zúñiga-Mayo, Gerardo Armas-Caballero, Alexander DeLuna, Nayelli Marsch-Martínez, Stefan de Folter

## Abstract

Flowers are composed of different organs, whose identity is defined at the molecular by the combinatorial activity of transcription factors (TFs). MADS-box TFs interact forming complexes that have been schematized in the quartet model. The gynoecium is the female reproductive part in the flower, crucial for plant reproduction, and fruit and seed production. Once carpel identity is established, a gynoecium containing many tissues arises. Several TFs have been identified as regulators of gynoecium development, and some of these TFs form complexes. However, broad knowledge about the interactions among these TFs is still scarce. In this work, we used a systems biology approach to understand the formation of a complex reproductive unit as the gynoecium by mapping binary interactions between well-characterized TFs. We analyzed over 3500 combinations and detected more than 200 protein-protein interactions (PPIs), resulting in a process specific interaction map. Topological analyses suggest hidden functions and novel roles for many TFs. Furthermore, a relationship between TFs involved in auxin and cytokinin signaling pathways and other TFs was observed. We analyzed the network by combining PPI data, expression and genetic data, allowing us to dissect it into several dynamic spatio-temporal sub-networks related to gynoecium development subprocesses.

## INTRODUCTION

Fruits and seeds are the basis of the food supply. Fruit and flower development in Arabidopsis has been studied for more than 20 years (Roeder and Yanofsky, 2006; Alvarez-Buylla et al., 2010). The accepted model for floral organ formation relies on the combinatorial action of mostly MADS-box proteins, resulting in the specification of the four whorls of the flower: sepals, petals, stamens and carpels (Coen and Meyerowitz, 1991). At the molecular level, MADS-box transcription factors (TFs) form complexes in order to regulate gene expression. For the female reproductive organs, the C-class and E-class proteins physically interact to give the identity to the gynoecium (Honma and Goto, 2001; Theissen and Saedler, 2001).

Gynoecium formation starts at stage 6 of flower development, when its identity is established (Smyth et al., 1990). After that, several TFs participate in the subsequent formation of the tissues. These TFs belong to a significant number of TF families such as MADS-box, HOMEOBOX, bHLH, Zinc Finger, bZIP, factors mediating hormone transcriptional responses (ARF, ARR, TCP), and others (Roeder and Yanofsky, 2006; Reyes-Olalde et al., 2013).

Current regulatory networks underlying gynoecium patterning include many protein-encoding genes, mostly TFs, acting in a coordinated fashion for the formation of all the tissues (Chavez-Montes et al., 2015; Marsch-Martinez and de Folter, 2016; Zúñiga-Mayo et al., 2019). The involvement of several TFs in the same process generates robustness. At the very early stages, the establishment of two domains, the lateral and the medial, is essential for further gynoecium development. Besides TFs, the hormones auxin and cytokinin are involved (Bowman et al., 1999; Sehra and Franks., 2015; Marsch-Martinez and de Folter, 2016; Muller et al., 2017; Victor-Zuñiga et al., 2019).

Subsequently, the establishment of the carpel margin meristem or CMM is critical (Wynn et al., 2011; Reyes-Olalde et al., 2013). All the medial tissues will be formed from the CMM, including the placenta and ovules, septum, transmitting tract, replum, style and stigma. During these early developmental processes, important TFs are SHOOT MERISTEMLESS (STM), CUP-SHAPED COTYLEDON (CUCs), SPATULA (SPT), ETTIN (ETT), and some B-type ARABIDOPSIS RESPONSE REGULATORS (ARRs) (Sessions et al., 1997; Heisler et al., 2001; Nahar et al., 2012; Kamiuchi et al., 2014; Scofield et al., 2007; Reyes-Olalde et al., 2017).

After that, in the formation of the transmitting tract another set of TFs are involved such as NO TRANSMITTING TRACT (NTT) (Crawford et al., 2007), the HECATE proteins (HEC1, HEC2 and HEC3), with SPT (Gremski et al., 2007), HALF FILLED (HAF), BRASSINOSTEROID ENHANCED EXPRESSION 1 and 3 (BEE1, BEE3), and the AUXIN RESPONSE FACTORS 6 and 8 (ARF6, ARF8) (Crawford and Yanofsky, 2011). In the medial-lateral patterning of the gynoecium, this is, the formation of valves, valve margins, and replum, a repression-action between lateral and medial factors is necessary, ASYMMETRIC LEAVES 1 and 2 (AS1, AS2), JAGGED (JAG), FILAMENTOUS FLOWER (FIL), YABBY 3 (YAB3), and FRUITFULL (FUL) in the lateral domain; on the other hand, NTT (Marsch-Martinez et al., 2014), BREVIPEDICELLUS (BP), SHOOT MERISTEMLESS (STM) and REPLUMLESS (RPL) (Roeder et al., 2003), promote replum formation in the medial domain (Alonso-Cantabrana et al., 2007; Gonzalez-Reig et al., 2012); while, in between those tissues, SHATTERPROOF 1 and 2 (SHP1, SHP2), ALCATRAZ (ALC), INDEHISCENT (IND), and SPT are involved in the formation of the valve margins at later stages of gynoecium formation (Roeder and Yanofsky, 2006; Girin et al., 2011).

Although there is information about how the above-mentioned TFs perform their activities, most of that information has come from studies done at the individual level. However, virtually all TFs form protein complexes (Vidal et al., 2011; Herrera-Ubaldo et al., 2014; Bemer et al., 2017). For all the TF families listed above, some essential feature of the protein functioning is achieved via PPIs, although, probably we know only the tip of the iceberg. Therefore, generating more PPI data, i.e. direct binary interaction mapping in a high-throughput, standardized manner, would boost our understanding on TF functioning during plant development.

As expected, also for gynoecium and fruit development, information on PPIs is rather scarce (Herrera-Ubaldo et al., 2014). Studies have been reported on the generation of PPI data for some TF families such as MADS-box (de Folter et al., 2005; Immink et al., 2009), Homeobox (Hackbusch et al., 2005), TCP (Danisman et al., 2013), and proteins involved in hormone response like ARF-Aux/IAA (Vernoux et al., 2011) or ARRs (Dortay et al., 2006). In other cases, information comes from the study of a specific tissue or a specific developmental process, e.g. transmitting tract (Gremski et al., 2007) or fruit elongation (Ripoll et al., 2015). Some putative TF complexes have been predicted, such as SEU-LUG-FIL-ANT for the control of the adaxial fate in the gynoecium (Azhakanandam et al., 2008), and ETT-IND-BP-RPL (Simonini et al., 2018) for style development. In summary, current information about physical interactions important for gynoecium and fruit development is the result of very focused studies.

In this work, we made a systematic effort to boost the information on PPI data that can give us insights into gynoecium and fruit development from the perspective of PPI networks of TFs. The formation of the gynoecium is a complex process, where many actors participate, so the study of this complex process as a system is necessary. We generated a PPI map of TFs that are expressed in the gynoecium and/or fruit and tissues with meristematic activity, and combined this with functional information and the sequence of developmental events that occur to obtain a proper gynoecium and fruit. This is an important step towards a comprehensive gynoecium gene regulatory network, and, furthermore, will be a useful resource for understanding how TFs can function. A deep understanding of molecular mechanisms and the regulatory networks controlling fruit development could be a tool for facing the future problems in crops and food production related to climate change (Zhao et al., 2017).

## RESULTS

Here we present a protein-protein interaction network (PPI network) with many of the well-characterized transcription factors (TFs) involved in gynoecium development in *Arabidopsis thaliana* (hereafter Arabidopsis). The combination of functional information and the physical interaction data allowed us to extract networks most likely reflecting protein complexes directing the formation of tissues, and with the addition of manually curated expression data we studied the dynamics of interactions related to temporal cues. With these, we dissected the interaction map into 13 sub-networks, each network representing a region and a progressive time during gynoecium development. These sub-networks are likely biologically relevant.

### Transcription factors involved in gynoecium development physically interact

The first question we wanted to address was, how the TFs involved in gynoecium development physically interact between them. We selected well-studied TFs (and some co-factors and other TFs) related to this process (most of them are listed in Reyes-Olalde et al., 2013), in total, our study included 72 proteins (gyn set; listed in Supplemental Table 1). We performed a matrix-based yeast two-hybrid (Y2H) assay with Arabidopsis protein-encoding ORF clones. To date, this is still the technique that has generated most PPI data for many model organisms, in a high-throughput, standardized and reproducible manner (Walhout and Vidal, 2001; Vidal et al., 2011; Braun et al., 2013). Most of the possible combinations were tested, but some of the BD clones were not used due to high autoactivation (results of the autoactivation tests are shown in Supplemental Table 2). In total, 3648 combinations were tested (Supplemental Table 2). For the Y2H assay, three interaction reporters were used; two of them allowing yeast growth (-HIS and -ADE), and colonies were confirmed with a third colorimetric reporter (LacZ). The interaction dependent growth was monitored with four yeast crosses (biological replicates), with one technical replicate, giving a total of 8 colonies per marker; one replicate from each marker was used for the LacZ assay, adding four more score points.

**Table 01.** Topological comparison of the 13 sub-networks.

| Zone |  | Nodes | Edges | Clustering coefficient | Diameter | Radius | Network centralization | Shortest paths | Char. path length | Avg. number of neighbors | Density | Heterogeneity | Isolated nodes | Self-loops |
| --- | --- | --- | --- | --- | --- | --- | --- | --- | --- | --- | --- | --- | --- | --- |
| 1 | Gynoecium primordium | 46 | 162 | 0.34 | 5 | 1 | 0.40 | 1894 | 2.226 | 6.61 | 0.147 | 0.867 | 0 | 10 |
| 2 | Lateral domain | 10 | 4 | 0.00 | 2 | 1 | 0.19 | 8 | 1.25 | 0.60 | 0.067 | 1.106 | 5 | 1 |
| 3 | Medial domain | 30 | 43 | 0.21 | 7 | 4 | 0.24 | 552 | 2.953 | 2.47 | 0.085 | 0.986 | 6 | 6 |
| 4 | Valves | 14 | 17 | 0.09 | 3 | 2 | 0.35 | 90 | 1.844 | 2.14 | 0.165 | 0.786 | 4 | 2 |
| 5 | Young replum | 6 | 6 | 0.44 | 3 | 2 | 0.30 | 20 | 1.5 | 2.00 | 0.4 | 0.577 | 1 | 0 |
| 6 | Carpel margin meristem | 33 | 58 | 0.19 | 7 | 4 | 0.23 | 756 | 2.86 | 3.09 | 0.097 | 0.947 | 5 | 7 |
| 7 | Replum | 11 | 22 | 0.50 | 3 | 2 | 0.39 | 72 | 1.444 | 3.82 | 0.382 | 0.589 | 2 | 1 |
| 8 | Medial region | 25 | 48 | 0.35 | 5 | 3 | 0.48 | 462 | 2.247 | 3.36 | 0.14 | 0.959 | 3 | 6 |
| 9 | Ovules | 32 | 72 | 0.18 | 5 | 3 | 0.38 | 812 | 2.296 | 4.06 | 0.131 | 0.911 | 3 | 7 |
| 10 | Stigma | 8 | 10 | 0.44 | 2 | 1 | 0.43 | 20 | 1.3 | 1.75 | 0.25 | 0.892 | 3 | 3 |
| 11 | Style | 12 | 12 | 0.18 | 3 | 2 | 0.36 | 72 | 2.028 | 1.67 | 0.152 | 0.894 | 3 | 2 |
| 12 | Septum | 13 | 20 | 0.39 | 3 | 2 | 0.36 | 56 | 1.571 | 2.31 | 0.192 | 0.952 | 5 | 5 |
| 13 | Transmitting tract | 6 | 10 | 0.67 | 1 | 1 | 0.30 | 12 | 1 | 2.00 | 0.4 | 0.707 | 2 | 4 |

In summary, each combination was tested 24 times. Plates were incubated during six days at 22°C. Colony growth was scored on the 6th day after inoculation. In total, yeast growth was detected in 294 combinations. For the construction of the network and further analysis we used the top scored interactions, these are, interactions detected with at least 2 out of the 3 reporters, with at least half of the colonies; this list includes 239 of high confident interactions (all the interactions are listed in Supplemental Table 3), which were used to generate the network. Almost one-third of the interactions had the highest score (24 points). In some cases, due to autoactivation of the BD clone, just two reporters could be used; in these cases, the highest score was 12 points (70 interactions). After removing duplicated edges, the network has 239 interactions (edges) between 58 proteins (nodes) (Figure 1 and Supplemental Figure 1).

**Figure 1.**
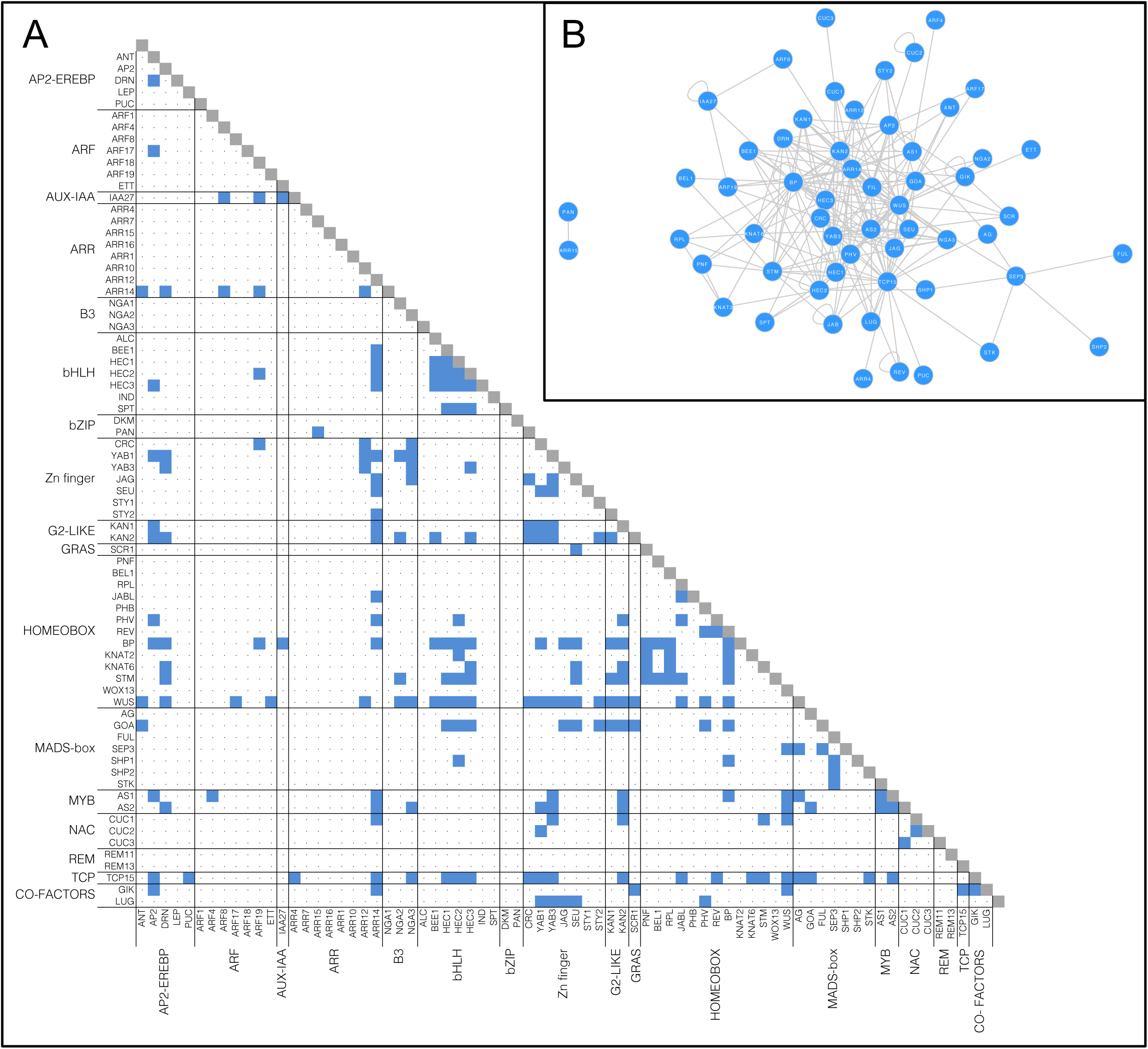
Transcription factors (TFs) involved in gynoecium development interact physically. (A) Matrix representing the interactions detected in the Y2H assay, proteins are grouped in families. (B) Protein-protein (PPI) network constructed with the information in (A).

We detected one major network containing 56 nodes, in addition to the interaction PAN-ARR15 that is not connected to the main network. A basic topological analysis was performed with the major network. Nodes with the highest number of connections (degree) and the value of other topological indexes are listed in Supplemental Tables 4 and 5. Interestingly, PPIs are not restricted to members of the same family of TFs (Figure 1A), and members of the HOMEOBOX and Zinc-finger family could interact with members of all the TF families we tested.

### Proteins involved in the same developmental processes are related at the physical interaction level

The PPI map we generated shows a single major component with practically all the interactions, so we wondered if TFs grouped according to their functions (i.e., the formation of a tissue). Since most of the TFs selected for this study have been previously characterized, we could add a tag related to their function or based on the phenotypes observed in the mutants, i.e., determinacy, style formation, etc. (Supplemental Table 1). With these, we generated networks highlighting proteins participating in 10 different developmental processes (Figure 2, Supplemental Figure 2). In general, at the level of physical interactions, proteins involved in the same processes do not cluster together in the network (Figure 2A-C, Supplemental Figure 2), but when those nodes are extracted from the network, we can see sub-networks or so-called modules, probably representing protein complexes (Figure 2D-F, Supplemental Figure 2).

**Figure 2.**
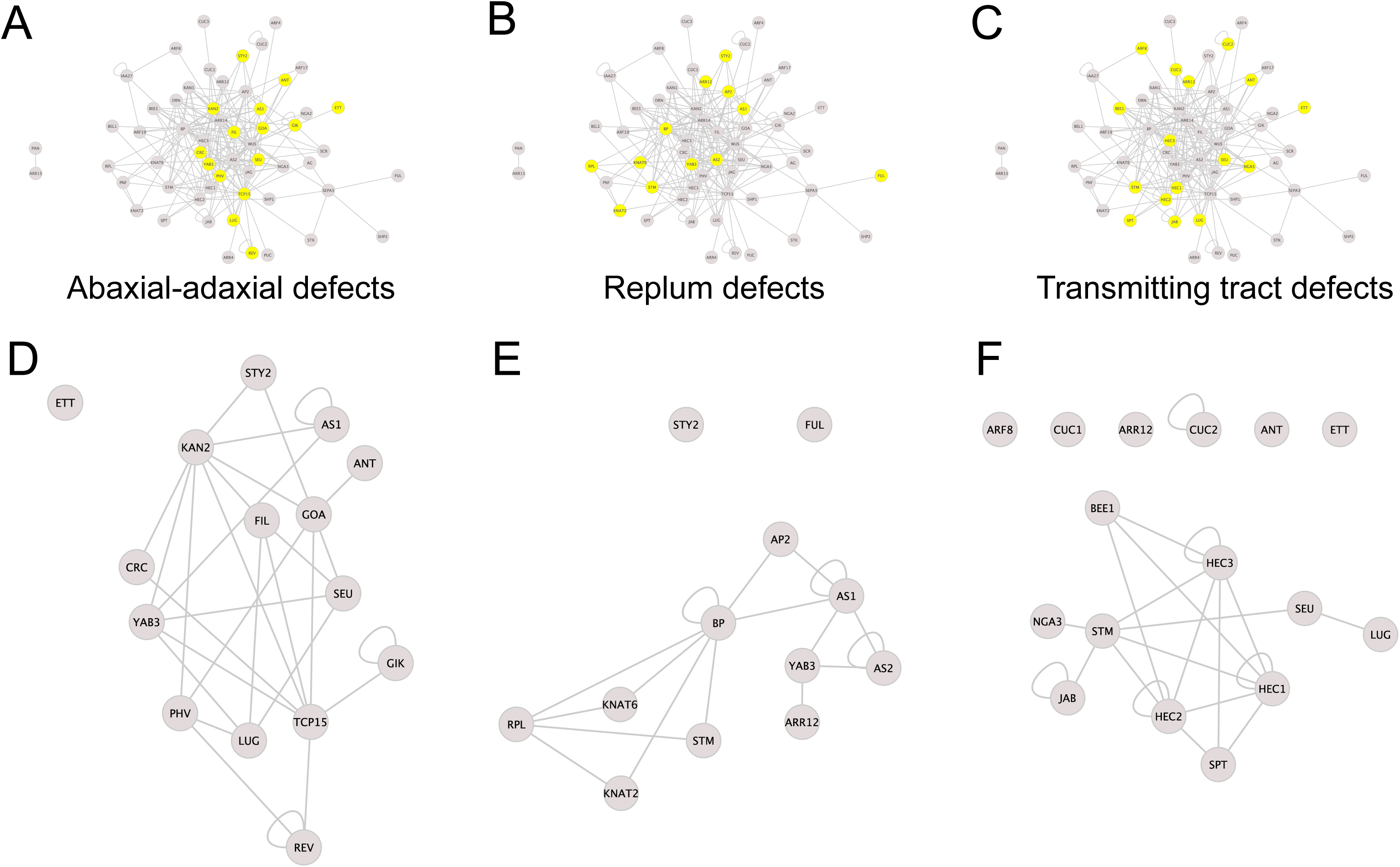
Proteins involved in the same process interact physically. (A-C) Representation of processes comprising gynoecium development within the interaction network, the proteins involved are colored in yellow (D-F).

With the tested interactions, we observe that sub-networks related to carpel number establishment or determinacy are not as complex (number of nodes and interactions) as those related to ovule or abaxial-adaxial patterning. On the other hand, the networks corresponding to valve margins and replum are very similar.

### Dynamic interactions during gynoecium development

The sub-networks generated with the separation of interactions according to a functional process is an indicator of the complexity of the processes. It is likely that all the interactions do not occur at the same time. To address the temporal issue, and for a better visualization of gynoecium development through time, we generated a graphic outline of this process (Figure 3). Its construction is based on the progressive formation of tissues over time (Larsson et al., 2014; Reyes-Olalde et al., 2013), each region delimits a tissue or group of cells that can be defined as different (Figure 3A). This scheme uses lines as a representation of the different zones within the gynoecium; at the very beginning, the gynoecium primordium is represented as a single line, shortly after is divided in 2 (lateral and medial domains, visible at floral stage 7), and after that many other lines arise (floral stages 8 to 11), as a representation of all the tissues composing a mature gynoecium observed at floral stage 12. With this, we defined more than 10 spatio-temporal contexts (or zones) in which different protein interactions could be taking place during gynoecium development (Figure 3B).

**Figure 3.**
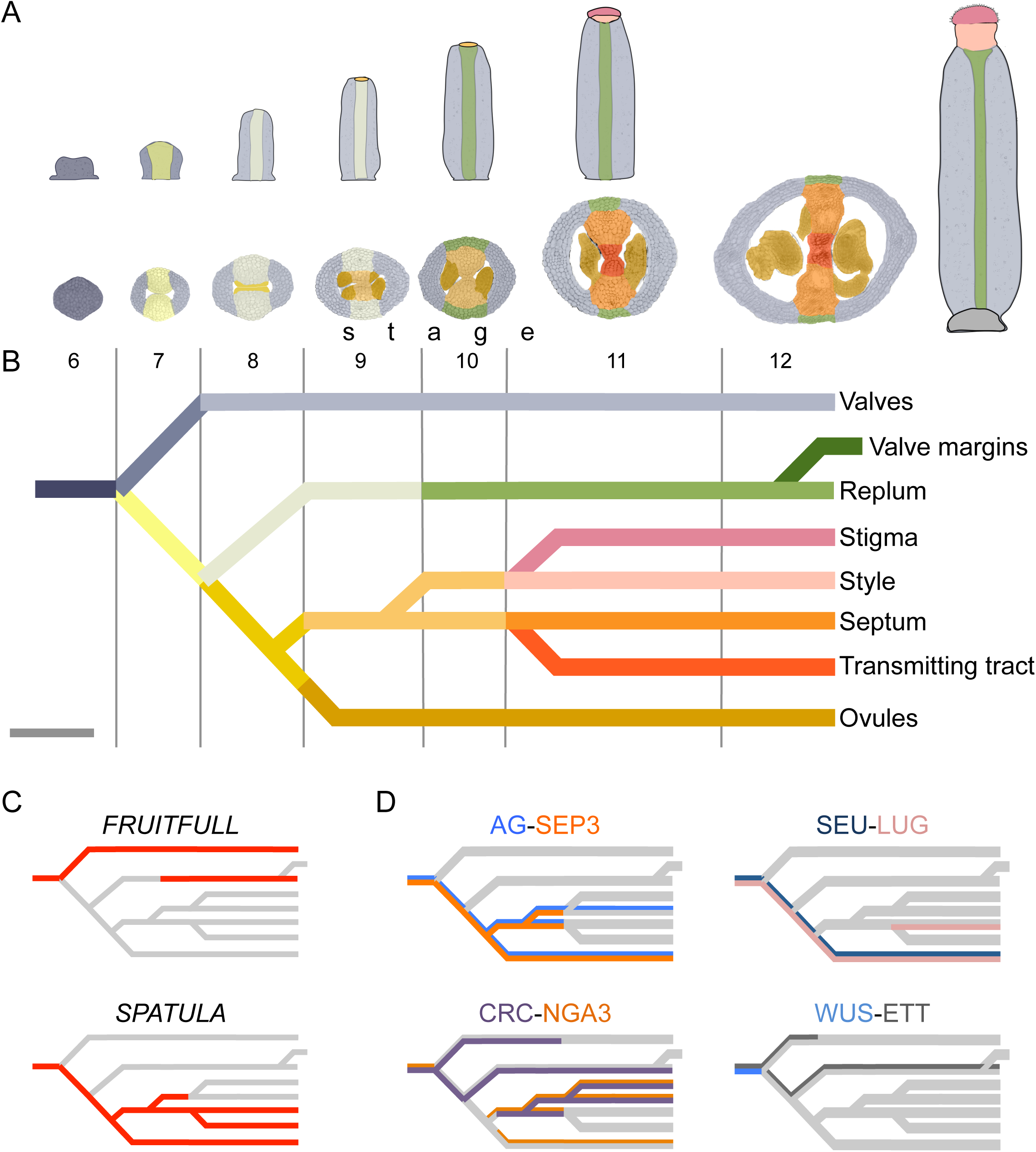
Gynoecium development over time. (A) Stages of gynoecium development, false colors represent different tissues within the gynoecium. (B) Schematic representation of gynoecium development, lines represent regions depicted in (A); scale bar represents 1 day. (C) Expression patterns of *FRUITFULL* and *SPATULA*, lines represent those regions indicated in (B). (D) Interaction patterns of the dimers AG-SEP3, SEU-LUG, CRC-NGA3 and WUS-ETT, colored lines represent the expression pattern of the protein-encoding gene in the same color.

This scheme helped us to address questions such as where and when the identified interactions are likely to occur during gynoecium development. We took advantage of all the previous studies performed for each gene; most of them at the individual level, a large proportion of this data was previously collected (Reyes-Olalde et al., 2013). In this study we updated the information, added data for other genes, manually curated gene expression data and represented it according to our division of tissues (listed in Supplemental Table 1).

This graphical representation is useful for displaying individual gene expression patterns during gynoecium development (Figure 3C). The overlapping of expression patterns with proteins that physically interact is an approach to study interaction profiles of two proteins, revealing where and when that dimer may be performing its functions (Figure 3D). In some cases, near perfect matches in expression pattern coincidence are observed, such as AG-SEP3 and SEU-LUG; in other cases, as WUS-ETT or CRC-NGA3 partial matches in expression patterns suggest that those interactions could occur in specific regions and stages of the gynoecium, and interactions change if different contexts.

The use of expression data allowed to dissect the PPI network in a spatio-temporal manner; nodes representing TFs were placed in each of the contexts we defined, so we can have a PPI network for each context (Figure 4, Supplemental Figure 3), these networks represent the subset of interactions that could be occurring in those tissues at specific times. The networks are different in node and edge number; the differences in complexity between them can be easily observed (Table 1). A large fraction of the complete network is observed at the beginning of gynoecium formation. After that, the lateral domain recruits a small part and most of the nodes and interactions are restricted to the medial domain. This network remains very similar for the carpel margin meristem (CMM), ovules and septum; and near floral stage 10 is separated into 4 networks, representing style, stigma, transmitting tract and septum. The networks for valves and replum are similar, they start as small networks, and some nodes and interactions appear over time.

**Figure 4.**
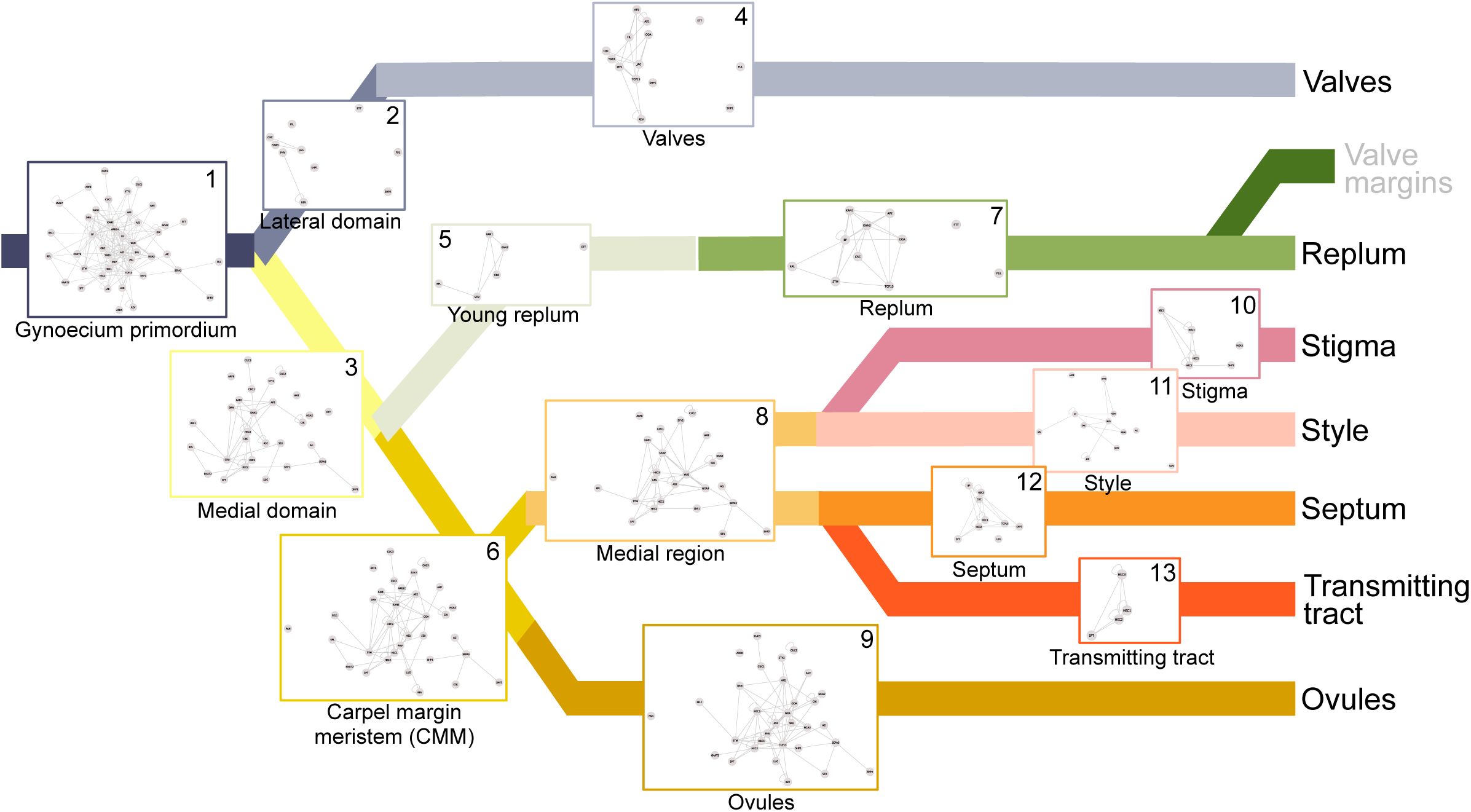
Succession of protein interaction networks during gynoecium development. The combination of gene expression data and physical protein-protein interaction (PPI) information allowed the dissection of the network into 13 sub-networks underlying the formation of tissues or developmental sub-processes.

### TF interactions and hormonal pathways

Transcriptional responses to hormones (i.e., auxin and cytokinin) are mediated by a set of response regulators: Auxin Response Factors (ARFs) and Arabidopsis Response Regulators (ARR, A and B-types) (Schaller et al., 2015). Some of these proteins were included in our interaction test. We detected interactions between the hormone response regulators and TFs from various families (Figure 5A). Interestingly, five TFs, WUS, HEC2, CRC, AS1, and BP can physically interact with regulators of both hormonal signaling pathways, suggesting that these nodes may serve as links to connect hormonal pathways.

**Figure 5.**
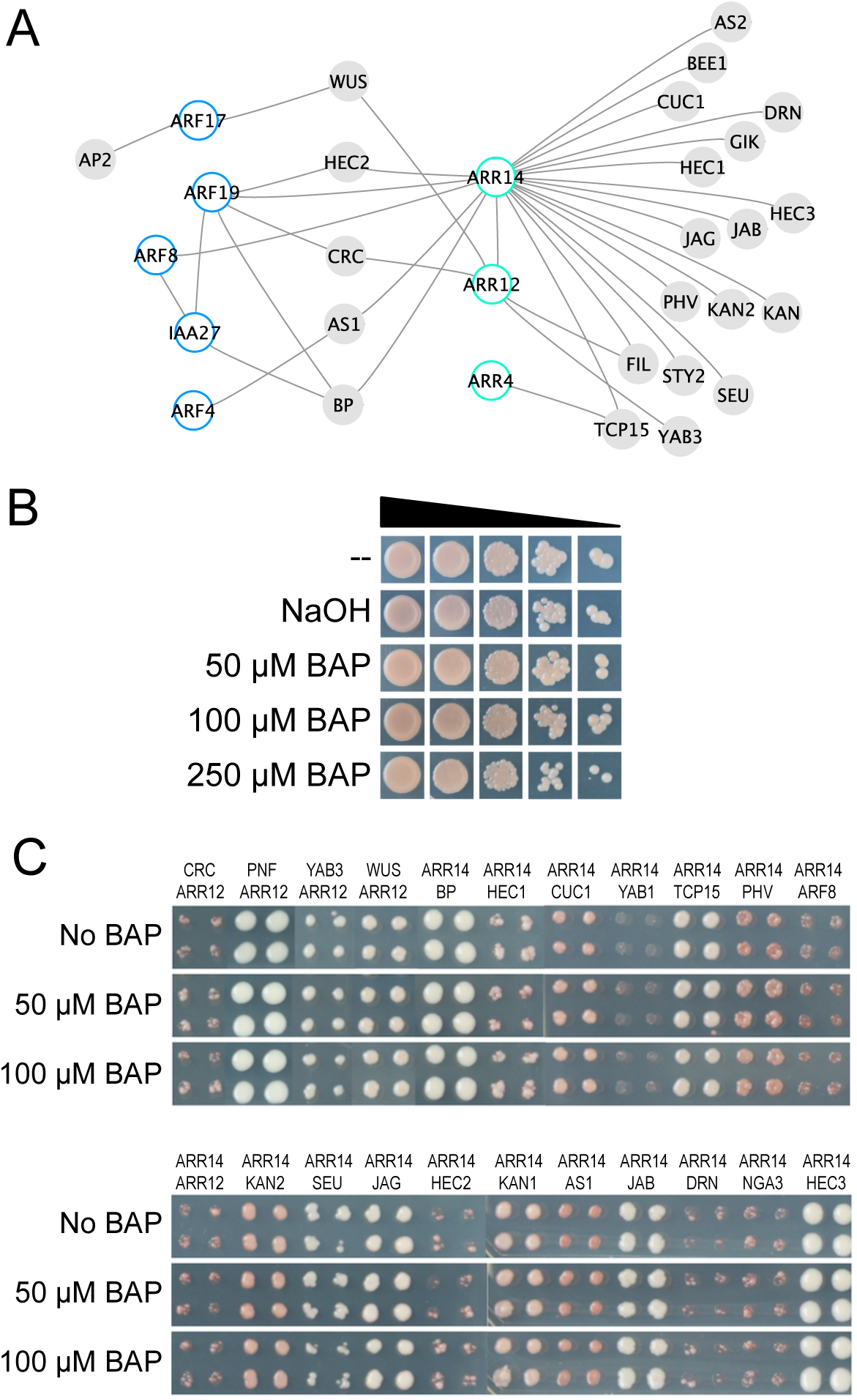
The effect of cytokinin on protein interactions. (A) Sub-network including transcription factors involved in auxin and cytokinin signaling pathways and their interactions with other proteins. (B) The addition of BAP to the culture medium does not affect the growth of the PJ69 yeast strain, neither when diluted yeast concentrations are inoculated (indicated with the triangle; dilution series 1x, 10x, 100x, 1000x, and 10000x). (C) The addition of BAP to the culture medium does not affect the interaction-dependent growth of yeast; some of the combinations shown in (A) are presented on medium containing 0 μM, 50 μM, or 100 μM BAP.

Another interesting phenomenon in the context of protein interactions and hormones is a recent report showing that the interaction-dependent yeast growth could be modified due to auxin addition to the culture medium (Simonini et al., 2016). In order to test if this phenomenon is occurring also with our set of interactions, we used the colonies recovered (representing a dimer or an interaction) from the -ADE experiments (171 interactions, listed in Supplemental Table 3). We used the -ADE set because the entire set of interactions could be tested using the same medium (test with –HIS would involve the addition of different concentrations of 3-AT). First, we tested the effect of two hormones on the wild type PJ69 yeast strain. The addition of cytokinin (BAP: 6-Benzylaminopurine) did not affect the growth of the yeast, even at the maximum tested concentration (250 μM) (Figure 5B). On the other hand, the effect of the addition of auxin (IAA: Indole-3-Acetic Acid) on the yeast growth is observed in the dilution series; the highest concentration of inoculum seems not to change yeast growth at 20, 50 or 100 uM IAA, but is affected when 250 uM IAA was tested (Supplemental Figure 5). For the Y2H test, we inoculated the colonies in hormone-containing media at different concentrations, and incubated them at 22°C during 6 days. In our conditions, no significant changes in yeast growth were detected due to cytokinin addition (Figure 5C), or auxin addition (Supplemental Figure 5).

### The extended PPI network and novel actors of gynoecium development

The PPI network we generated includes well-characterized TFs controlling gynoecium development (72 protein-encoding genes we selected for this study, gyn set, Figure 6A), so it can be considered as a process specific interaction map. Our map contains 239 interactions between 58 proteins (Figure 6B, listed in Supplemental Table 6). For some proteins we did not detect interactions in our system, which can be due to various reasons (e.g., missing TFs in our gyn set, weak interaction strength, formation of higher-order complex needed, etc.)

**Figure 6.**
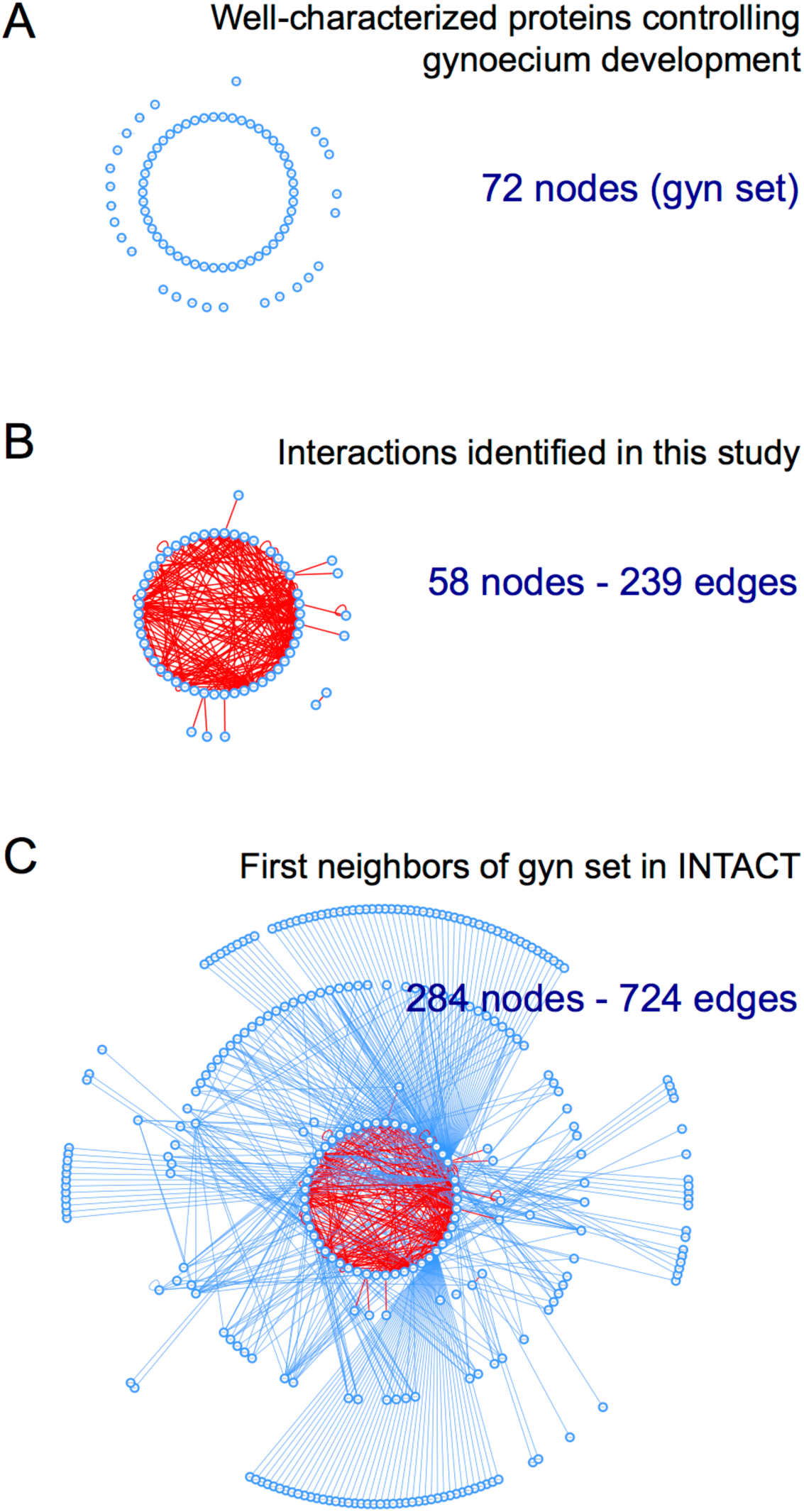
The extended network of transcription factors controlling gynoecium development in Arabidopsis. (A) Number of proteins used in this study, the gyn set. (B) PPI data identified for the gyn set (this work). (C) The extended network, the PPI data of this work combined with PPI data from the IntAct database (based on Y2H for our gyn set nodes and only of protein-encoding genes with expression in carpels).

So, to partially overcome this and identify novel TF interactors, we generated an extended PPI network (Figure 6C). The extended PPI network includes reported interactions (IntAct database; Orchard et al., 2014) between the gyn set of proteins (this study) and other TFs, most of the times uncharacterized proteins and obtained in studies not focused on gynoecium development. In this extended network, we included only interactions detected by Y2H assays and furthermore, this list was filtered to include just TF-encoding genes expressed (at least) in carpels (Supplemental Table 6; purple colored selection, Supplemental Figure 6). This extended network can be considered as a comprehensive interaction network of TFs involved in gynoecium development (724 interactions, Figure 6C). All the interactions in the extended network are listed in the Supplemental Table 6. Because our gyn set consists out of well-studied TFs, in the extended network, for every dimer in the list at least one partner has been characterized, meaning we can predict a role for the other TF. In conclusion, this paves the way for focused functional studies of over 200 TFs present in the extended network as candidates for having a role in the regulation of gynoecium development. We present a list of 26 TFs (of the extended PPI network) that have the highest number of interactions (> 4) with the TFs of the gyn set (see Supplemental Table 6). In general, these TFs share various interactors, and, notably, in the top 10 of TFs with most interactions, there are 9 TFs of the TCP family.

### An Aux/IAA TF affects reproductive development

One of the proteins found to interact with known gynoecium TFs in the extended PPI network is IAA27, an Aux/IAA protein not reported as regulator of Arabidopsis gynoecium development, but with potential roles in this process based on the interactions found (Figure 6C, Supplemental Table 5). We obtained the AD and BD clones and included it in the Y2H assay, where reported interactions were confirmed, and novel interactions were detected (Figure 1 and Supplemental Figure 5). Subsequently, to study the potential function of IAA27, wild type Arabidopsis plants were transformed with an antisense construct to knock-down the expression of this gene (*anti-IAA27*). As presumed, different phenotypes were observed during the reproductive phase of *anti-IAA27* plants (Supplemental Figure 5). Expression analysis showed that the level of *IAA27* silencing was correlated with the severity of the observed phenotypes. Independent lines with a severe phenotype were characterized in more detail. The inflorescences of *anti-IAA27* plants produced fewer floral buds compared with wild type inflorescences. Furthermore, the floral organs showed a yellowish aspect, the anthers were shorter with a reduced number of mature pollen grains, and the short fruits produced on average only two seeds.

Different analyses were made to determine why *anti-IAA27* plants developed short fruits. Most of the ovules analyzed showed defects in their morphology, mainly an arrest in embryo sac development. Transverse sections showed that anthers of *anti-IAA27* plants produced fewer pollen grains compared with wild type anthers. Finally, the *LAT52::GUS* reporter line was used to visualize pollen tube growth. Gynoecia from wild type and *anti-IAA27* plants were hand-pollinated with *LAT52::GUS* pollen. Pollen tubes grew efficiency through wild type gynoecia; however, pollen tubes grew only in the style but not in the ovary of *anti-IAA27* gynoecia (Supplemental Figure 5). These results indicate that both male and female reproductive development is affected in *anti-IAA27* plants. Consistent with this, reproductive development is also affected when the function of its interactor ARF8 is abolished (Nagpal et al., 2005; Wu et al., 2006). Furthermore, it was recently shown that silencing of the homologue of *IAA27* (*SlIAA27*) in tomato also affects male and female reproduction (Bassa et al., 2012).

## DISCUSSION

In the past years, gynoecium development has been studied in a gene-by-gene fashion. Current knowledge for gynoecium development includes dozens of genes and their genetic interactions, though information on protein-protein interactions (PPI) is available but still rather scarce. With the PPI map generated in this study, we want to add a solid basis for the molecular understanding and the generation of a more complete gynoecium GRN. Recently, we reported some steps towards a comprehensive and dynamic GRN for gynoecium formation (Chavez Montes et al., 2015). Furthermore, all the data presented here will be a useful resource for people studying reproductive development.

### A network for gynoecium development

With the interactions among TFs reported here, we have generated the most complete PPI map for gynoecium development in Arabidopsis to date. Sub-networks or modules were considered as basic units, or fundamental units guiding morphogenesis, as seen in biological networks (Alon, 2003; Zhu et al., 2007). We identified 239 interactions using the gyn set (Figure 1), placing our PPI network at the position of various other PPI networks in plants (Braun et al., 2013). In addition, as discussed below, we generated an extended PPI network for gynoecium development containing 724 interactions (Figure 6).

The yeast 2-hybrid (Y2H) system is one of the most used techniques for the identification of binary protein interactions, with the advantage of being high-throughput, highly standardized, and reproducible, especially when using ORF collections (Walhout and Vidal, 2001; Vidal et al., 2011; Braun et al., 2013). The most complete interaction network in Arabidopsis to date, has been built using the Y2H system (AI-2, Arabidopsis Interactome Map Consortium, 2012), with 5664 interactions between 2661 proteins. Several other networks for Arabidopsis have been constructed with this type of data e.g., ARF-Aux/IAA (Vernoux et al., 2011), TCS (Dortay et al., 2006 & 2008), TOPLESS proteins (Causier et al., 2012), protein-G complex (Klopffleisch et al., 2011), ESCRT proteins (Richardson et al., 2011), cell cycle protein (Boruc et al., 2010), TCP family (Danisman et al., 2013), and MADS-box proteins (de Folter et al., 2005), among others.

The construction of a PPI map guiding gynoecium and fruit development and the GRNs underlying the process is a starting point for the study of the formation of the tissues, comparison of fruit development between species, the study of network rewiring allowing the formation of novel structures or processes in a global way. It can be used for the study of the evolutionary history of this important reproductive unit.

### Emerging properties for well-studied transcription factors

The network we generated, has two components, with the exception of the dimer PAN-ARR15, all the nodes form a major network component (Figure 1). This is an indicator of the close relationship between all the proteins and the processes involved in gynoecium development. Topological analysis is useful to get insights in network architecture and performance (Zhu et al, 2007). Analysis of the main component indicates that it is a compact network (network diameter =5; radius =1; characteristic path length =2.26) (Supplemental Table 5). We compared our network to other plant PPI networks, and its density is similar to these other networks (density =0.13) (Supplemental Table 5). Furthermore, most of the other analyzed topological characteristics are also in the range of the reported biological plant PPI networks, supporting our findings. One interesting aspect is that the nodes are very different in connections (network heterogeneity =0.91), in the case of TFs involved in gynoecium development this heterogeneity is observed in the effect of mutations on the genes, since not all cause the same defect.

Some topological indexes (see Supplemental Table 4), revealed hidden features for well-known TFs exposing interesting properties/activities for them, which will be an exciting topic for *in planta* and functional analysis. For instance, the high value in node degree (i.e. many interactions) for WUS, BP, STM, YAB1, HEC2 and HEC3, many involved in meristematic activity and all of them involved in early events of gynoecium development (Reyes-Olalde et al., 2013), and the network generated for this early stage (Figure 4, Table 1), is a sign of how complex this early event of gynoecium formation is.

Furthermore, the node TCP15 has also a high degree; this property is maybe the cause of the severe defects that occur in gynoecium development when *TCP15* it ectopically expressed (Lucero et al., 2015; Kieffer et al., 2011). Another similar example is ARR14 (B-type ARR) (Kieber and Schaller, 2014), a node with many interactions, suggesting essential roles in female reproductive development, though functional and expression data are still scarce, but the PPI data suggests a central role in gynoecium development.

The nodes WUS, TCP15, BP, and ARR14 have high values for stress and betweenness centrality (Supplemental Table 4), indicating a possible role as a mediator of interactions and maybe for the formation of high-order complexes, interestingly, SEPALLATA3 (SEP3) has similar values. This crucial protein for flower development has been described as a bridge for MADS-box protein interactions (Immink et al., 2009).

The values for clustering coefficient highlight proteins such as SPT, BEE1, and HEC1, as members of a highly connected module, probably reflecting their involvement in transmitting tract formation as a protein complex (Heisler et al., 2001; Gremski et al., 2007; Crawford et al., 2011). Other proteins with high values of clustering coefficient are AS2 and NGA3, suggesting that probably they can form higher-order complexes.

Neighborhood connectivity is an attractive property of nodes, meaning having few interactors but those interactors function as a hub (having many interactors); some of these proteins are ETT, AINTEGUMENTA (ANT), STYLISH 2 (STY2), and ARR4. In the case of gynoecium development, this differential interaction (heterogeneity) is maybe the cause for the formation of one tissue. The critical interaction could be the difference in the case of very similar networks, for instance, those from the medial region and the tissues it produces.

### Biologically relevant protein interactions

The occurrence of PPIs among TFs was tested in a heterologous expression system, namely yeast. As mentioned before, the Y2H system has many advantages and is an attractive technique for binary interaction mapping, which will continue (Vidal and Fields, 2014). As with these studies, next steps will be necessary to explore the functional relevance of detected interactions. The identification of physical interactions between proteins is just a starting point for functional studies and for understanding important developmental and cellular processes. Of course, starting with more focused functional analysis, various complementary techniques are available for PPI confirmation and to obtain further insight in e.g. protein complex formation, which are future perspectives of this work (Xing et al., 2016; Bontinck et al., 2018; Lampugnani et al., 2018).

Nevertheless, based on coexpression of the protein-encoding genes, we consider that most of the interactions could be biologically relevant because for 87% of the dimers, the genes are coexpressed (190 out of 219; based on the sum of the sub-networks in Supplemental Figure 3). This number is still an underestimation when we would consider e.g. more specific cell-type expression data, protein stability, and protein movement. Furthermore, another kind of evidence supporting the physical interactions detected here, is the interaction of proteins participating in the same biological process.

### Extended Network, a directed search of novel players in gynoecium development

The PPI network we generated in this work includes most of the well-characterized TFs involved in gynoecium development, though, it is highly likely that more TFs are involved. A rough estimation sets the expression of more than 500 TFs during carpel formation (de Folter at al., 2004; Chavez Montes et al., 2015). So, it is a challenge to characterize all of them. We made a first step to identify more TFs by exploring PPIs reported for our gyn set of TFs (IntAct database; Orchard et al., 2014). Many non-characterized proteins interact with the TFs of our network, however, we included only those TFs expressed in flower and fruit, which resulted in the extended PPI network with 724 interactions between 284 nodes (Figure 6). This approach proved to be valid since we validated at least one of them, the Aux/IAA IAA27 protein (Supplemental Figure 5). We found similar functions as one of its interactors ARF8, which is involved in reproductive development (Nagpal et al., 2005; Wu et al., 2006). Furthermore, an example of another biologically relevant PPI present in the extended network is between NO TRANSMITTING TRACT (NTT) and SEEDSTICK (STK), which we recently reported as important for medial domain development and plant reproductive competence (Herrera-Ubaldo et al., 2018).

### A combinatorial model for gynoecium development?

The quartet model is an elegant representation of the molecular mechanism underlying floral organ specification (Theissen and Saedler, 2001). One of our goals here was the proposal of a combinatorial model for gynoecium formation, an extension of the ABC model. In this sense, testing binary interactions between TF guiding gynoecium development is the first step towards a model explaining gynoecium patterning.

However, there are some crucial differences between the specification of the different floral organs in a flower versus the specification and formation of the different tissues in the gynoecium (i.e. after carpel specification), which, in general, is more complex: 1) The implication of several TF families (flower: MADS and AP2; and more than 16 TF families for gynoecium); 2) The number of TFs involved in the process (Flower <10; gynoecium >70), this is related to the redundancy observed between many of them; 3) The effect of mutations of the genes, for the flower, mutations in the homeotic genes produce dramatic changes in the architecture of the flower, while in gynoecium formation (i.e. after carpel specification), the mutations (often) generate less dramatic changes. 4) The time of the developmental processes, floral organ specification takes around four days; on the other hand, most of the tissues of the gynoecium (floral stages 6 to 12) are formed during more than ten days.

Considering those four main differences, a possible ‘model’ would involve a succession of networks, controlled in time. For flower formation, spatial organization is a key feature, observed with the spatially restricted interactions, while for gynoecium formation, both time and spatial organization are important during development.

### Inserting hormones into the networks

Hormones play a significant role during gynoecium development, particularly auxin and cytokinin (Marsch-Martinez and de Folter, 2016; Zuñiga-Mayo et al., 2019). Given the importance of these two hormones, we included various proteins involved in the transcriptional response into the network (ARFs and ARRs). There are other TFs that do not participate directly in the primary response to hormones, but their relationship with hormones has been demonstrated, e.g. SPT (Schuster et al., 2015; Reyes-Olalde et al., 2017), TCP15 (Lucero et al., 2015), HECs (Gaillochet et al., 2018), FUL (Ripoll et al., 2015), STM (Jasinski et al., 2005; Yanai et al., 2005), CRC (Yamaguchi et al., 2017), and AG (Ó’Maoiléidigh et al., 2018), among others.

Here, we found interactions between ARFs, ARRs, and TFs of different families. Especially, we detected five TFs (WUS, HEC2, CRC, AS1, and BP) that can interact with both auxin and cytokinin related TFs. These TFs are expressed in meristems and boundaries, so they may have a role as decision-makers, in the control of proliferation and differentiation. Other studies have observed similar cases (e.g. Oh et al., 2014; Yazaki et al., 2016; Gaillochet et al., 2018). Our detected PPIs give an opportunity to explain molecularly how these TFs can do so. It would be interesting to detect the dynamics of protein complex formation during different stages of gynoecium development, to know whether the identified interactions occur simultaneously or change in time.

Another interesting phenomenon involving PPIs and hormones is the ability of a hormone to modify the physical interaction of proteins, which has been observed recently for auxin with ETT (an ARF member) (Simonini et al., 2016), as well as affecting the transcriptional output (Simonini et al., 2017). Since we recovered all the colonies that grew in our Y2H experiments, we had the entire dimer collection in yeast colonies. So, we tested whether the gynoecium development PPI network could be modulated by the addition of auxin or cytokinin (IAA or BAP, respectively), however, in our conditions we did not detect alterations in PPI formation based on changes in yeast growth. Still this is an interesting topic to further explore, as well as directly in plants.

### The fruit is in the net

In this work, we started the study of gynoecium development from the systems biology perspective, the generation of a PPI map with many of the major players involved in this process, which gave us a vision of how complex the interplay could be between the TFs involved in the formation of tissues.

The next step, besides continuing with functional studies *in planta*, is to study the dynamical aspects (i.e. in spatio-temporal manner) of the interactions during gynoecium development. Moreover, TFs that could function as hubs in the network also deserve attention and should be further studied in this regard. Intriguing are the observed PPIs of TFs that interact with auxin and cytokinin related signaling proteins, and it will be very interesting to unravel the molecular meaning of this. Furthermore, we started studying the effects on PPIs by temporal cues and hormones. On the other hand, the protein complexes we proposed here, illustrated as sub-networks or modules need to be studied with complementary techniques such as bimolecular fluorescence complementation (BiFC) in different tissues *in planta* (Smaczniak et al., 2012; Long et al., 2017), co-immunoprecipitation (co-IP), protein arrays (Yazaki et al., 2016), and/or immuno or affinity purification followed by mass spectrometry analysis (IP-MS/AF-MS) (Smaczniak et al., 2012; Gadeyne et al., 2014; Huang et al., 2016).

We hope the information we presented here will serve as a resource and will be useful for other people studying plant reproductive development. Furthermore, since many TFs also participate in other developmental programs, some of these interactions may be conserved in other contexts, where they can be further explored.

## MATERIALS AND METHODS

### Clones

Information about the collection and generation of the Arabidopsis 72 ORF clones (the gyn set) is provided in Supplemental Table 2; many come from the REGIA collection (Paz-Ares and The REGIA Consortium, 2002). The gyn set is collection of mainly TFs that are expressed in the gynoecium and/or fruit and tissues with meristematic activity. Many of the well-characterized TFs involved in gynoecium development in *Arabidopsis thaliana* are present. Furthermore, a few cofactors are present as well.

### Yeast 2-hybrid assay

A matrix-based interaction assay was conducted as previously reported (de Folter and Immink, 2011). The GAL4 system (Invitrogen) was used; the coding sequences of the selected genes in entry clones were recombined with pDEST22 for GAL4-AD fusions or with pDEST32 for BD-fusions. Yeast transformation was performed with the PEG/LiAc method using the PJ69 yeast strain (James et al., 1996). All the tested combinations were generated by mating (see Supplemental Table 2), four AD colonies per construct were arranged in a single-well omnitray plate in a 384-colonies array. BD clones were grown individually in single-well plates. Mating was performed using a floating pin replicator (V&P scientific) in plates with YPAD medium. For diploid selection, colonies were transferred to plates with SD-TRP-LEU medium, the latter step was done twice. For the protein interaction test, diploid colonies were transferred to SD-LEU-TRP-ADE and to SD-LEU-TRP-HIS with different concentrations of 3-Amino triazole (see Supplemental Table 2), each selection marker was tested in duplicate. Plates were incubated at 22°C. Yeast growth was scored 6 days after inoculation. For the LacZ assay the RO-BLUE medium was used (de Folter and Immink, 2011) for these tests, one plate per protein was tested. Yeast colonies from both markers were recovered and cryo-conserved for future analysis (i.e., hormone dependent interactions).

### Hormone-dependent interaction assays

PJ69 yeast strain (James et al., 1996) and the colonies identified and conserved from the Y2H assay (-ADE set, Supplemental Table 3) were recovered and inoculated in medium SD-GLUC-TRP-LEU-ADE with increasing concentrations of cytokinin (6-Benzylaminopurine, BAP) or auxin (Indole-3-Acetic Acid, IAA). NaOH was used to help to dissolve the hormones to prepare stock solutions; 100 μM NaOH was tested as negative control for both, auxin and cytokinin experiments.

### Expression data

Data about gene expression for the selected protein-encoding genes were collected from literature; most of this information is an updated version of a previous report (Reyes-Olalde et al., 2013). For floral stage 6, RNA-seq data was collected from a previous report (Jiao and Meyerowitz, 2010), an expression value of >1.0 RPKM was considered. Separation in different regions during development was done according to previous works (Reyes-Olalde et al., 2013; Larsson et al., 2014). Functional processes were assigned according to the literature. All the information about expression and phenotypes is available in Supplemental Table 1.

### Interaction data collection and extended network construction

Protein interaction data was downloaded from IntAct (last accessed on 31.10.2018) (Orchard et al., 2014), the locus ID list of the 72 genes in the gyn set was used as query (Supplemental Table 1). Three filters were applied to the interaction list, 1) Interactions detected in Y2H assays (MI:2277, Cr-two hybrid; MI: 0726, Reverse two hybrid; MI:0018, two hybrid; MI:0397, two hybrid array; MI: 1112, two hybrid pooling approach; MI: 1356, validated two hybrid) were considered. 2) Interactions between transcription factors; a list was downloaded from the Plant Transcription Factors Database V4.0 (Jin et al., 2017) (last accessed 08.11.2018). 3) Expression in carpel; RNA-seq expression data was collected from ARAPORT11 (Krishnakumar et al., 2015) (last accessed 08.11.2018), genes with values of >0.5 TPM in carpel or stage 12 inflorescence samples were considered. Information about expression, GO classification and protein domains in the 221 TFs in the extended network (Supplemental Table 6 and Supplemental Figure 6) was collected from ARAPORT11 (Krishnakumar et al., 2015). The networks shown in Figure 6 are available in a Cytoscape network file (see Supplemental Figure 7).

### Network construction and topological analysis

Networks were generated using Cytoscape V3.6 (Shannon et al., 2003). Network analyses were performed with the Cytoscape built-in application NetworkAnalyzer (release 2.7; Assenov et al., 2008).

### Generation of *IAA27* knock-down lines and phenotypical characterization

A 293 bp fragment upstream of the stop codon of the *IAA27* gene (At4g29080) was PCR-amplified from cDNA using the primers 1KDL-IAA27 and 2KDL-IAA27 (see Supplemental Table 7), and subcloned into pENTR/D-TOPO (Invitrogen, Carlsbad, CA, USA). The pENTR/D-TOPO clone was sequence verified and recombined with the binary vector pHELLSGATE8 (Helliwell et al., 2002), resulting in the antisense construct *anti-IAA27.* The *anti-IAA27* lines were classified according to severity of fruit size phenotypes: strong if the fruit reached up to 50% of its normal size, medium if the fruit reached between 50 and 75% of its normal size, and weak if the fruit growth between 76-90% of its normal size.

For RT-PCR, plant tissue was collected for total RNA isolation according to the method described by Verwoerd et al. (1989). Total RNA was treated with DNaseI (Invitrogen) and reverse-transcribed using M-MLV (Invitrogen). Semiquantitative expression analysis for *IAA27* was performed using the primers 1RT-IAA27 and 2RT-IAA27 (see Supplemental Table 7). For pollination analysis, *LAT52::GUS* was used as pollen tube marker (Johnson et al., 2004). Gynoecia from *anti-IAA27* plants were emasculated 24 h before anthesis. The next day, anti-IAA27 gynoecia were hand-pollinated with pollen from *LAT52::GUS* marker line. 24 h after pollination, the gynoecia were collected and incubated 7 h at 37°C with a 5-bromo-4-chloro-3-indolyl-b-glucuronic acid solution (Gold Biotechnology, St Louis, MO, USA; Jefferson et al., 1987), placed in 90% ethanol for 1 h, followed by 70% ethanol for 1 h, and overnight in Hoyer solution (Anderson, 1954). Cross-section and GUS images were taken using a LEICA CTR6000 (Wetzlar, Germany) microscope in Differential Interference Contrast (DIC) mode.

## AUTHOR CONTRIBUTIONS

HHU and SdF conceived and designed the study; HHU, SEC, VLG, VMZM, and GAC performed the research; HHU and SdF analyzed the data; HHU performed visualization and prepared the figures; HHU, AD, NMM, and SdF discussed the research; HHU and SdF wrote the paper.

## ACKNOWLEDGMENTS

We would like to thank Jeanneth Pablo-Villa for initial assistance in the yeast 2-hybrid experiments in the lab. Furthermore, we the following people for clones: Cristina Ferrándiz, Alexander Heyl, Froukje van der Wal and Richard Immink, Eva Zanchetti and Lucia Colombo, Mitsuhiro Aida, Juan Carlos Ochoa-Sánchez, Irepan Reyes-Olalde, and the Arabidopsis Biological Resource Center (ABRC). We thank Karla González-Aguilera for technical assistance in the lab. HHU and VLG were supported by the Mexican National Council of Science and Technology (CONACyT) with a PhD fellowship (243380 and 487657, respectively). Work in the SdF laboratory was financed by the CONACyT grants CB-2012-177739 and FC-2015-2/1061, and NMM by the CONACyT grant CB-2015-255069. SdF acknowledges support of the Marcos Moshinsky Foundation and the European Union H2020-MSCA-RISE-2015 project ExpoSEED (grant no. 691109). No conflict of interest is declared.

## SUPPLEMENTAL INFORMATION

**Supplemental Table 1. List of genes selected for this study. Gene expression information and phenotype information. Related to Figures 1-4.**

**Supplemental Table 2. Yeast 2-hybrid experiment. List of clones, results of the autoactivation test, and tested combinations. Related to Figure 1.**

**Supplemental Table 3. Results of the Yeast 2-hybrid experiment. List of all the interactions detected, list of top-scored interactions used for the network construction, and list of interactions tested in the hormone assay. Related to Figure 1 and 5.**

**Supplemental Table 4. Topological analysis of the interaction map. Related to Figure 1.**

**Supplemental Table S5.** Topological comparison of plant interactomes. Related to Figure 1 and Table 1.

|  | A1 (a) | AUX signalling (b) | Cell cycle (c) | MADS (d) | Membrane (e) | Pathogens (f) | G-complex (g) | Gynoecium dev (h) |
| --- | --- | --- | --- | --- | --- | --- | --- | --- |
| Clustering coefficient | 0.0478 | 0.5574 | 0.1161 | 0.2228 | 0.1009 | 0.0239 | 0.0590 | 0.3122 |
| Connected components | 124 | 1 | 2 | 1 | 2 | 25 | 1 | 2 |
| Network diameter | 16 | 4 | 5 | 6 | 10 | 11 | 6 | 5 |
| Network radius | 1 | 2 | 1 | 3 | 1 | 1 | 4 | 1 |
| Network centralization | 0.0820 | 0.3682 | 0.2511 | 0.3604 | 0.1140 | 0.1084 | 0.3268 | 0.4016 |
| Shortest paths | 5767578 | 2256 | 1266 | 5550 | 35158 | 752650 | 187056 | 2972 |
| Characteristic path length | 4.749 | 1.726 | 2.657 | 2.637 | 4.226 | 4.786 | 3.326 | 2.259 |
| Avg. number of neighbors | 4.156 | 17.417 | 3.949 | 7.040 | 3.684 | 2.914 | 2.480 | 7.298 |
| Number of nodes | 2661 | 48 | 39 | 75 | 190 | 926 | 433 | 57 |
| Density | 0.0016 | 0.3706 | 0.1039 | 0.0951 | 0.0195 | 0.0031 | 0.0057 | 0.1303 |
| Heterogeneity | 2.336 | 0.578 | 0.683 | 0.867 | 1.170 | 2.307 | 3.676 | 0.906 |
| Isolated nodes | 27 | 0 | 0 | 0 | 0 | 0 | 0 | 0 |
| Self loops | 135 | 15 | 0 | 5 | 9 | 9 | 5 | 11 |
| Multi edge node pairs | 0 | 0 | 0 | 3 | 3 | 0 | 136 | 20 |

**Supplemental Table 6. Extended network. List of interactions in the extended network, list of proteins with >4 interactions, list of the 221 TF candidates for gynoecium development, and its classification in GO and protein domains. Related to Figure 6.**

**Supplemental Table S7.**
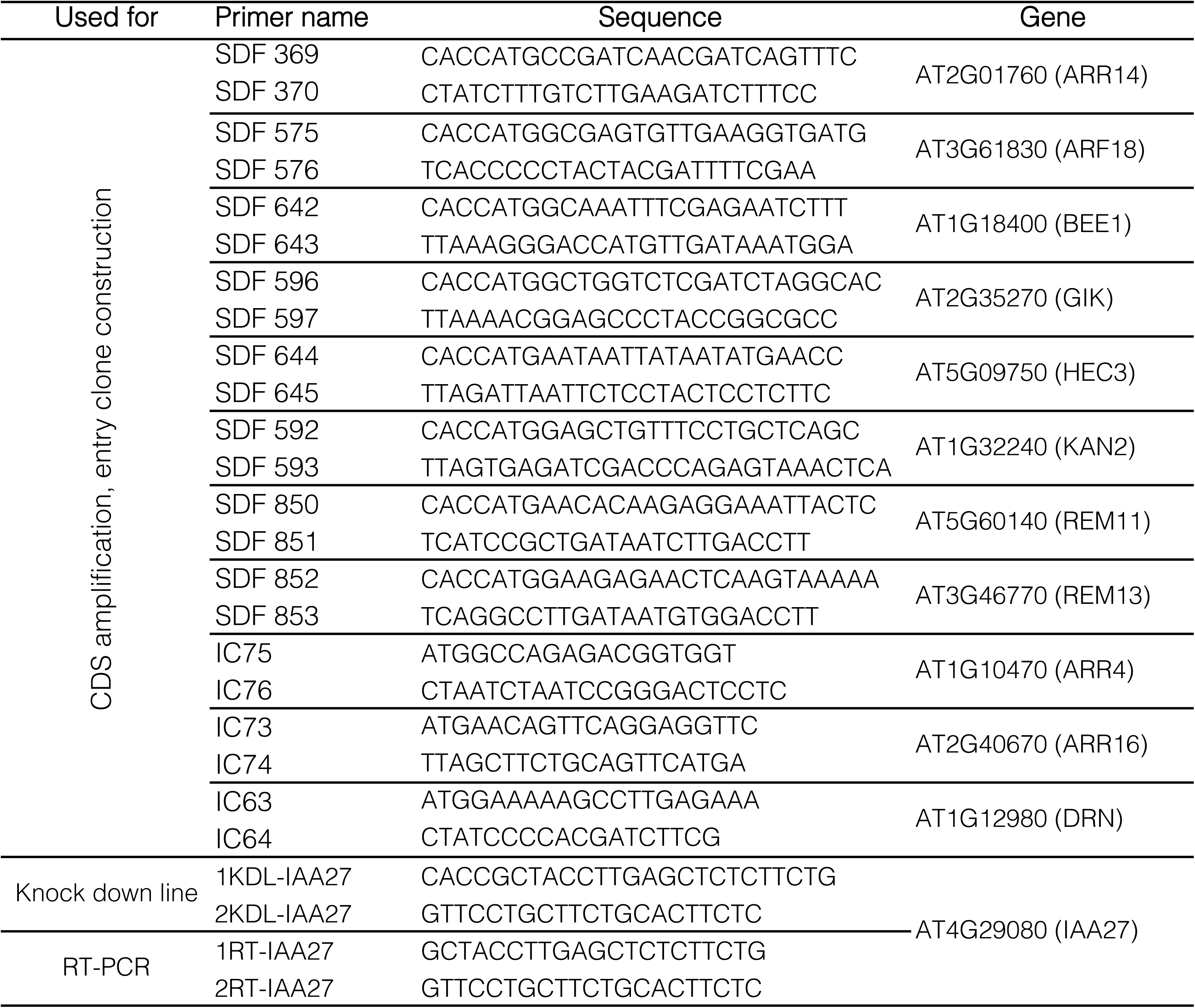
Primer list.

**Supplemental Figure 1.**
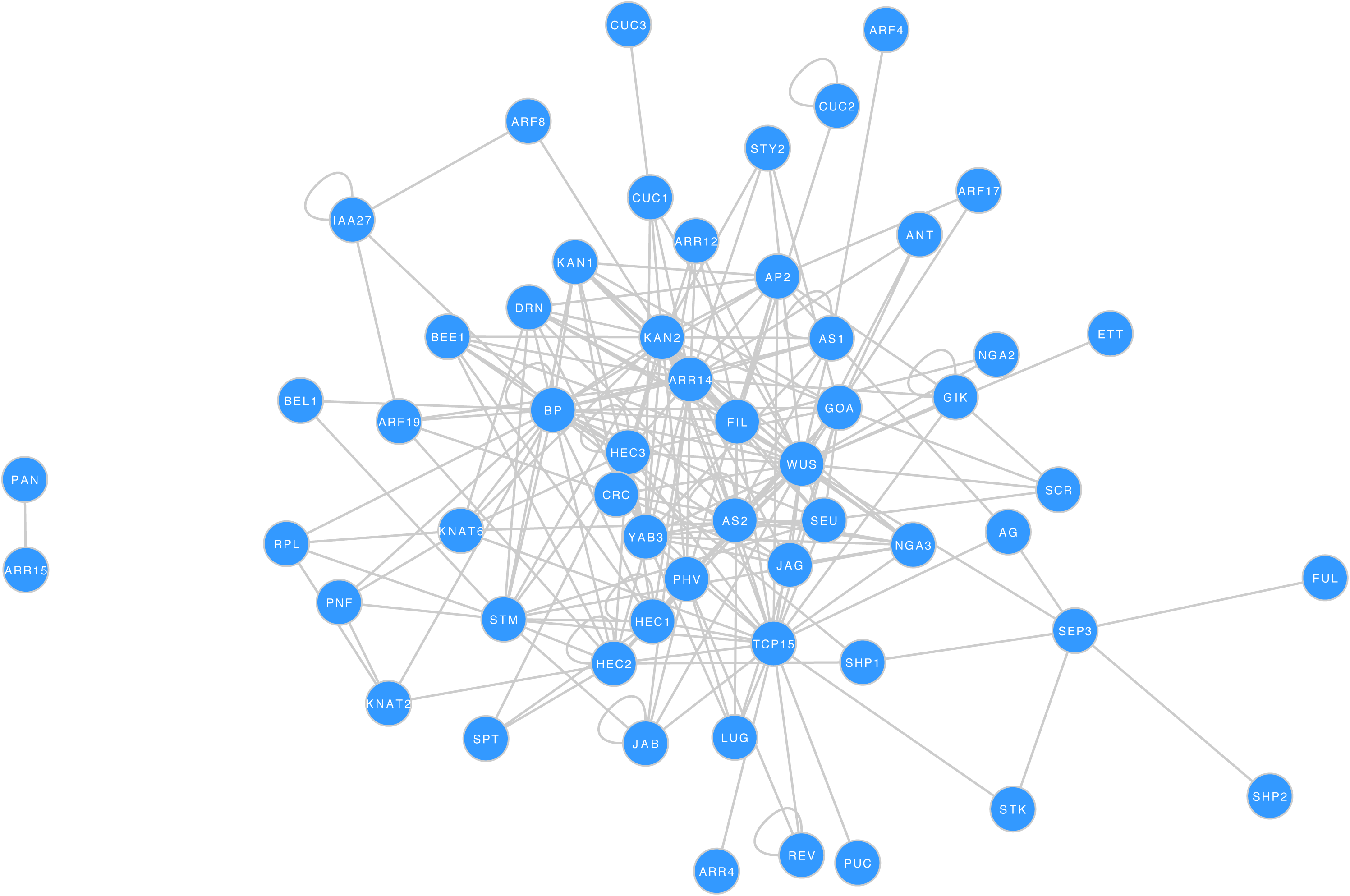
High-resolution protein interaction network of transcription factors controlling gynoecium development in Arabidopsis. Related to Figure 1.

**Supplemental Figure 2.**
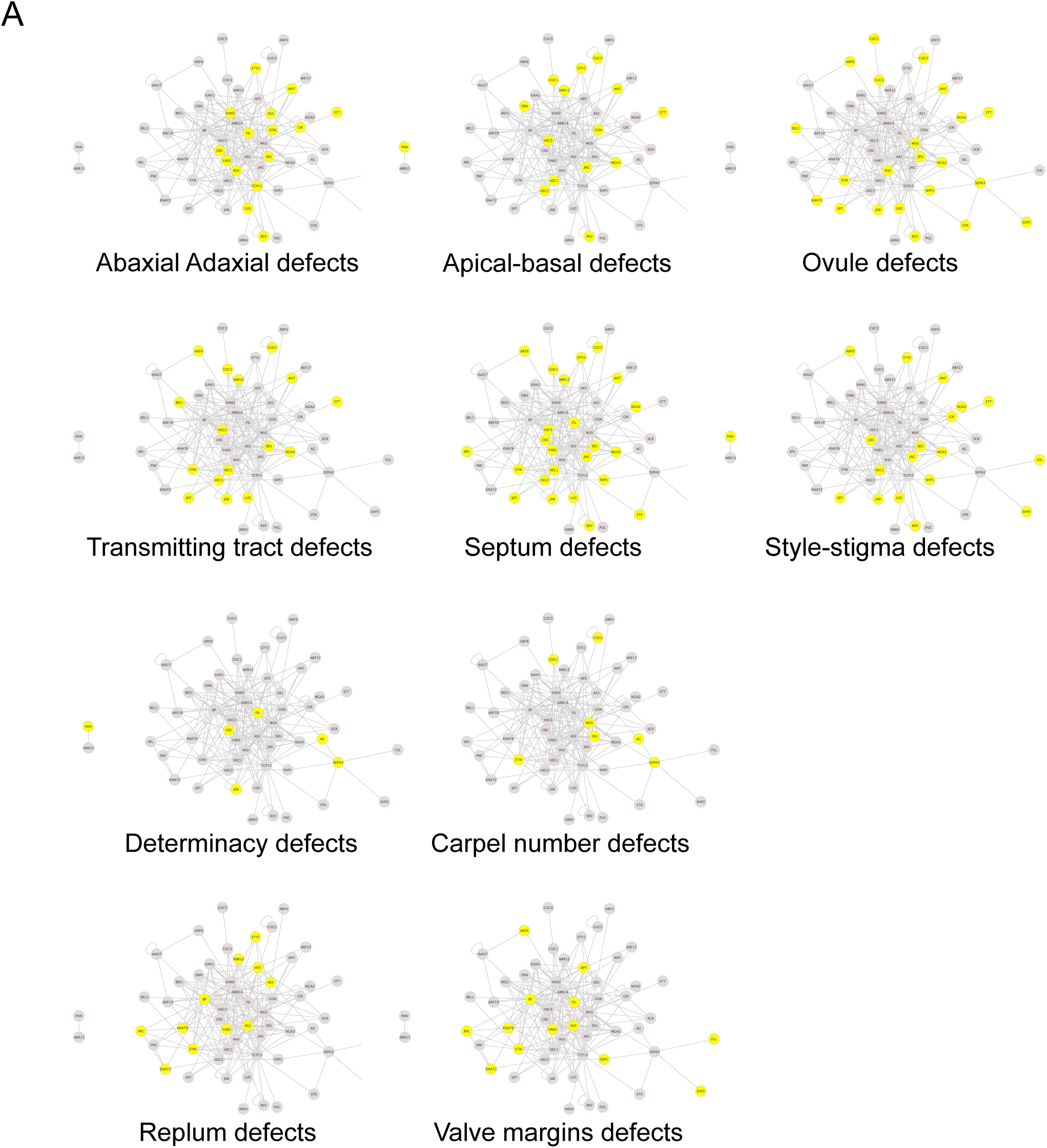
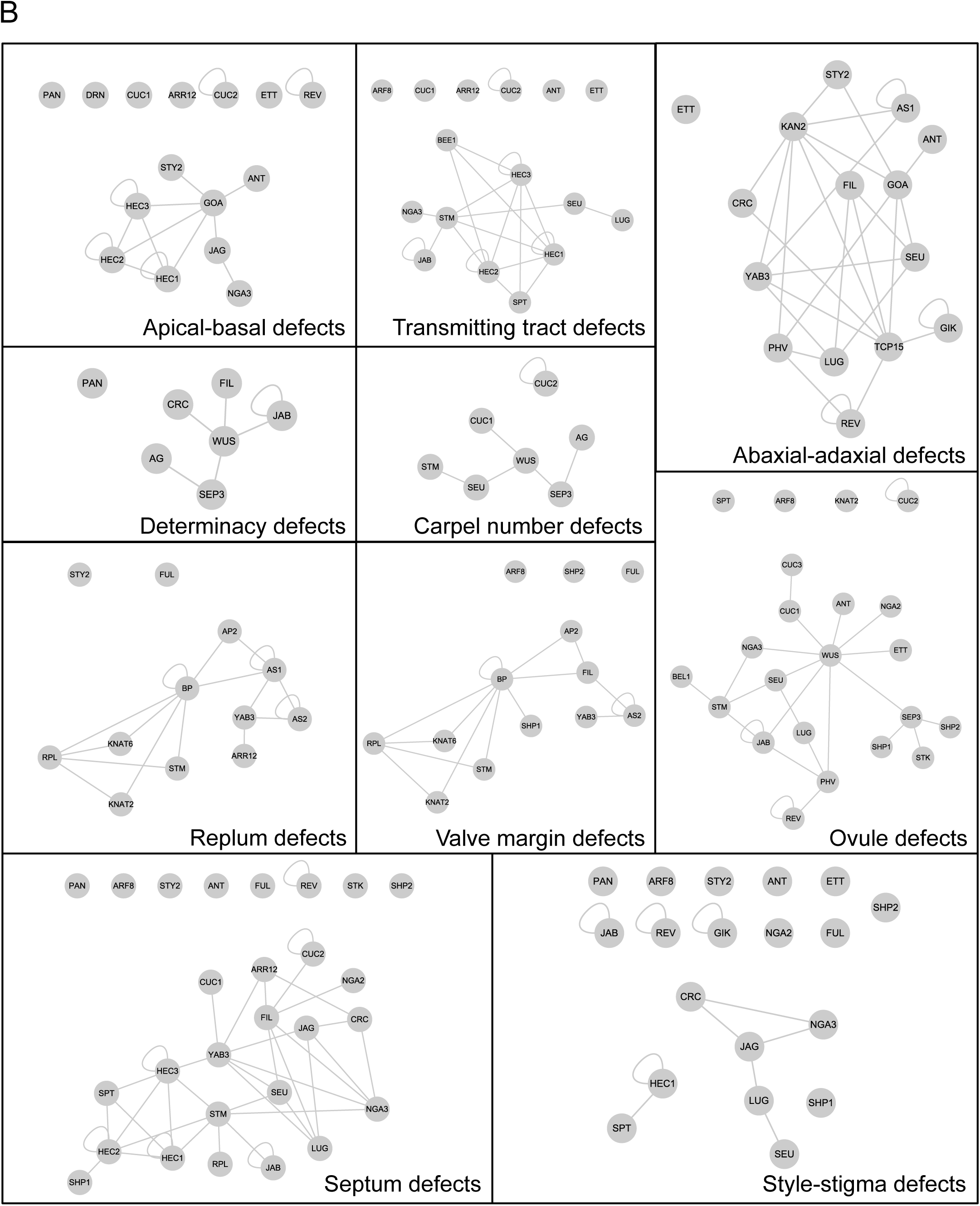
Networks related to phenotypes. Related to Figure 2. A) Nodes representing genes associated with a phenotype are selected in each network. B) Extraction of the selected nodes.

**Supplemental Figure 3.**
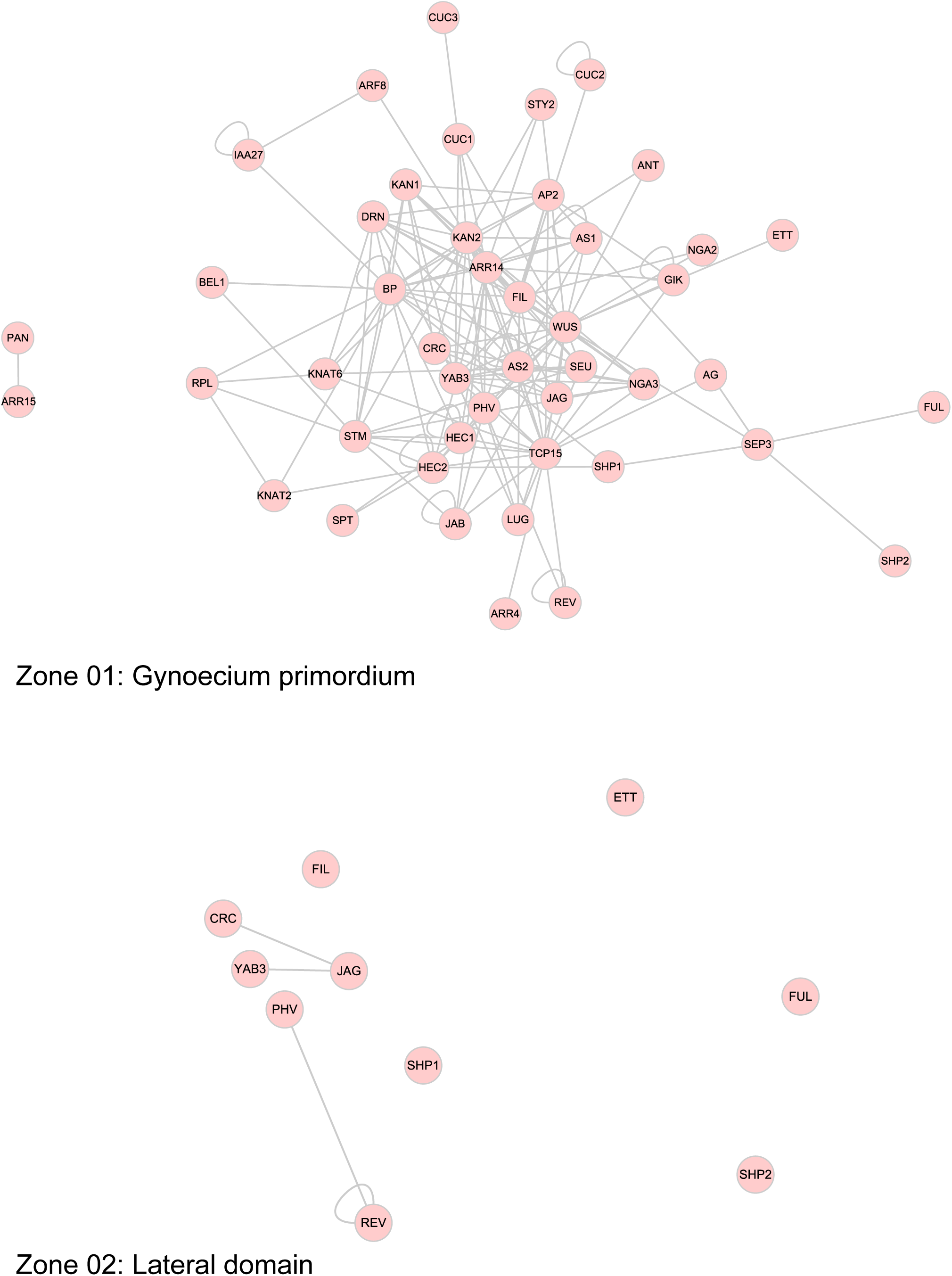
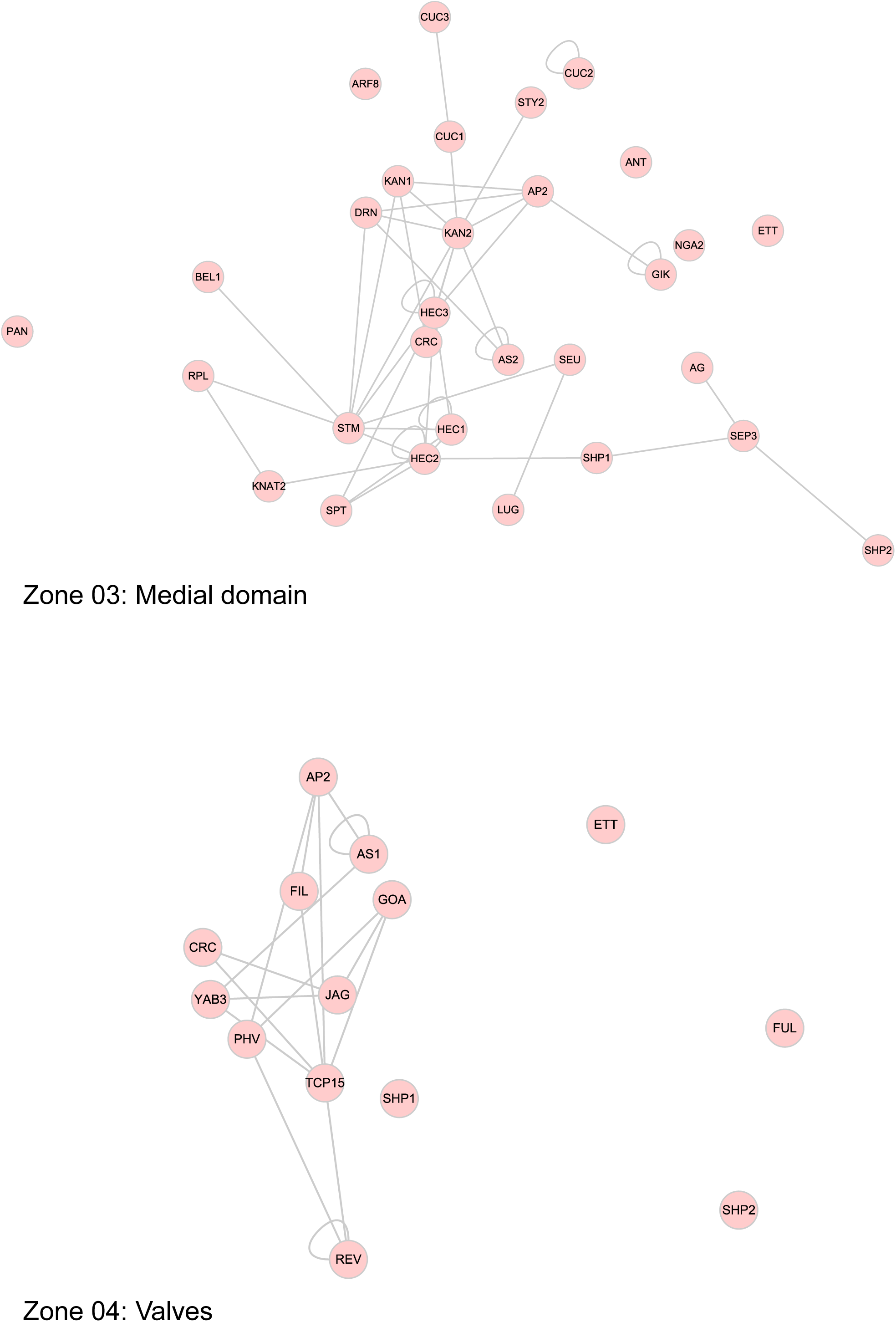
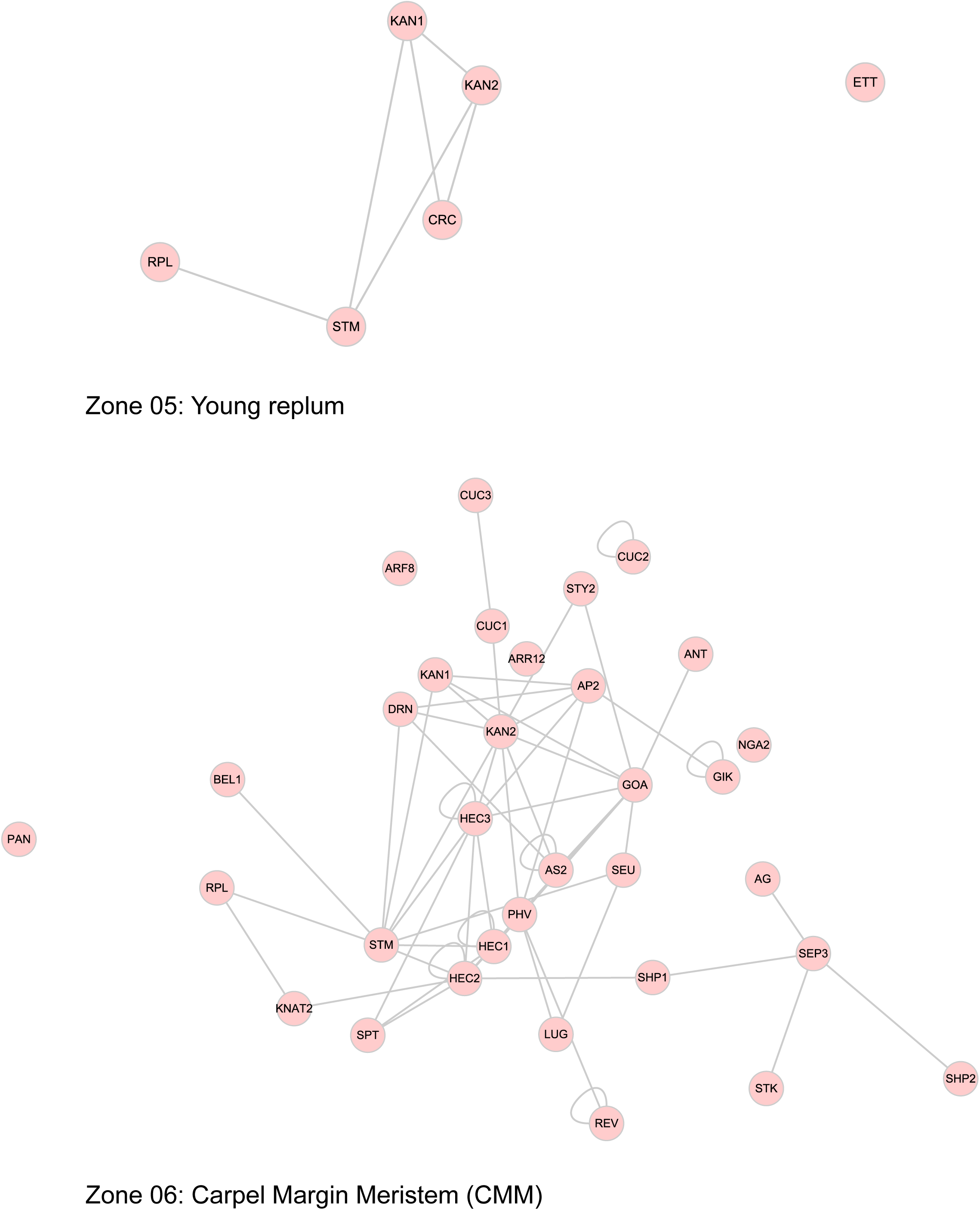
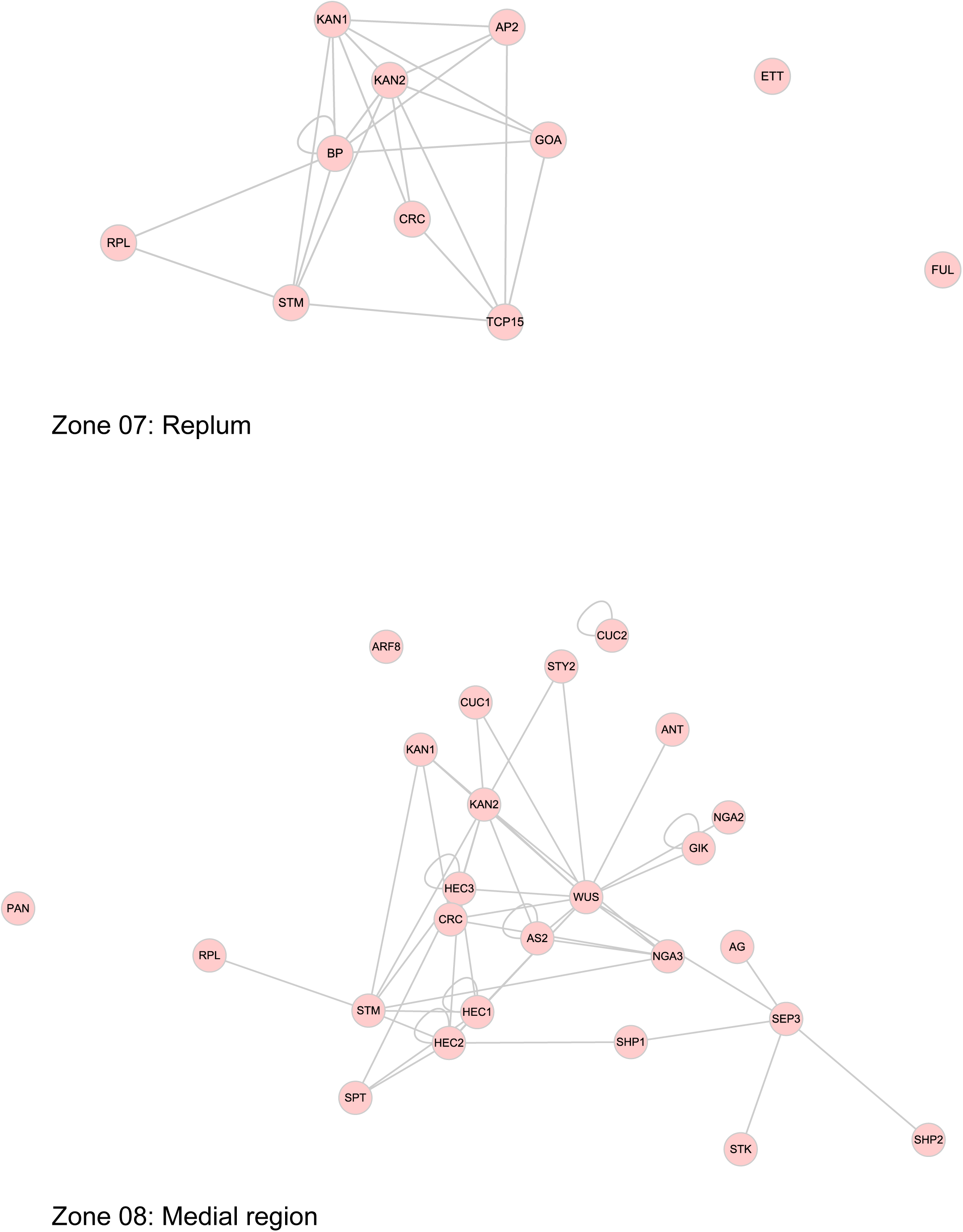
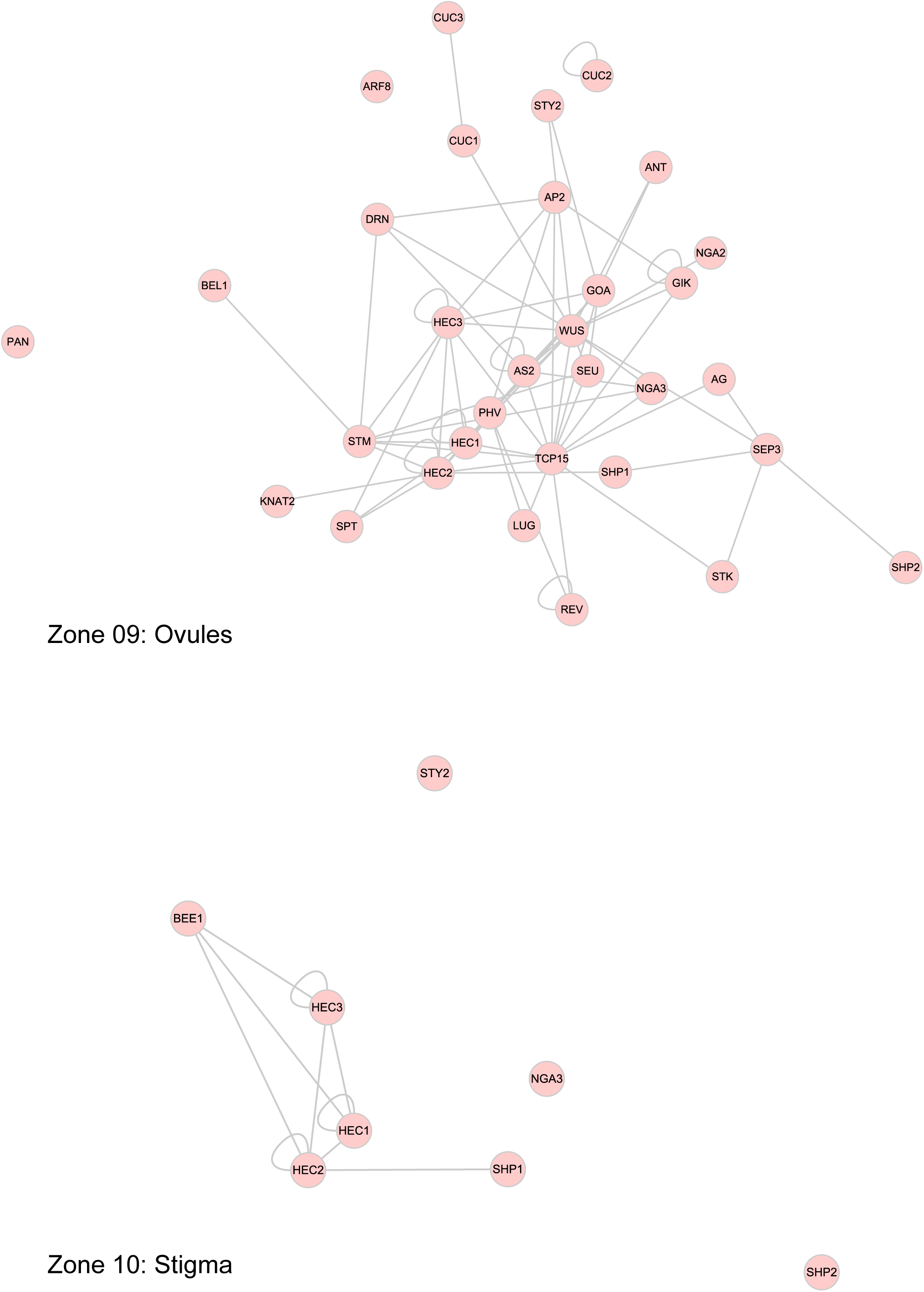
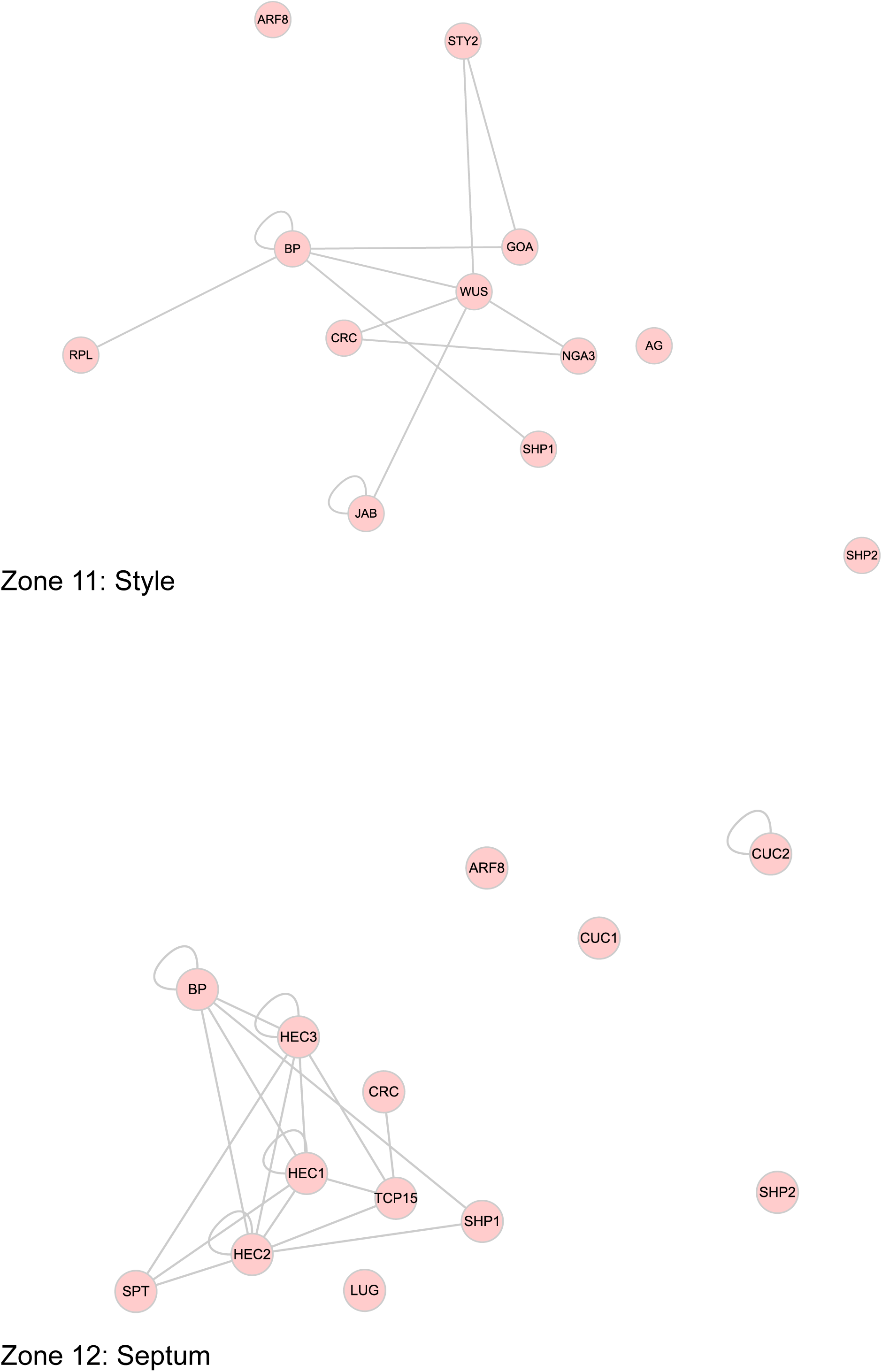
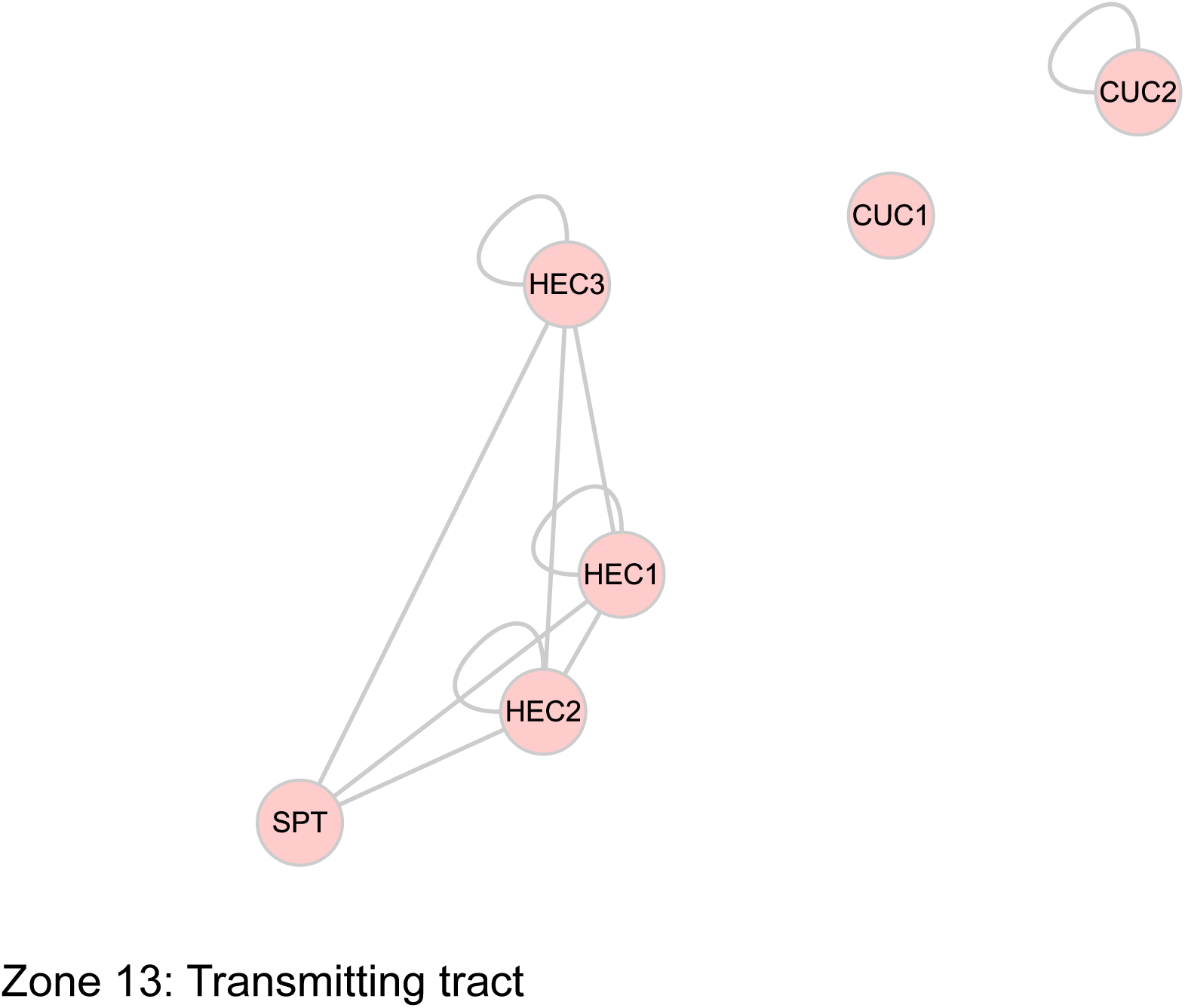
Networks for the 13 contexts (or zones) during gynoecium development. Related to Figures 3-4.

**Supplemental Figure 4.**
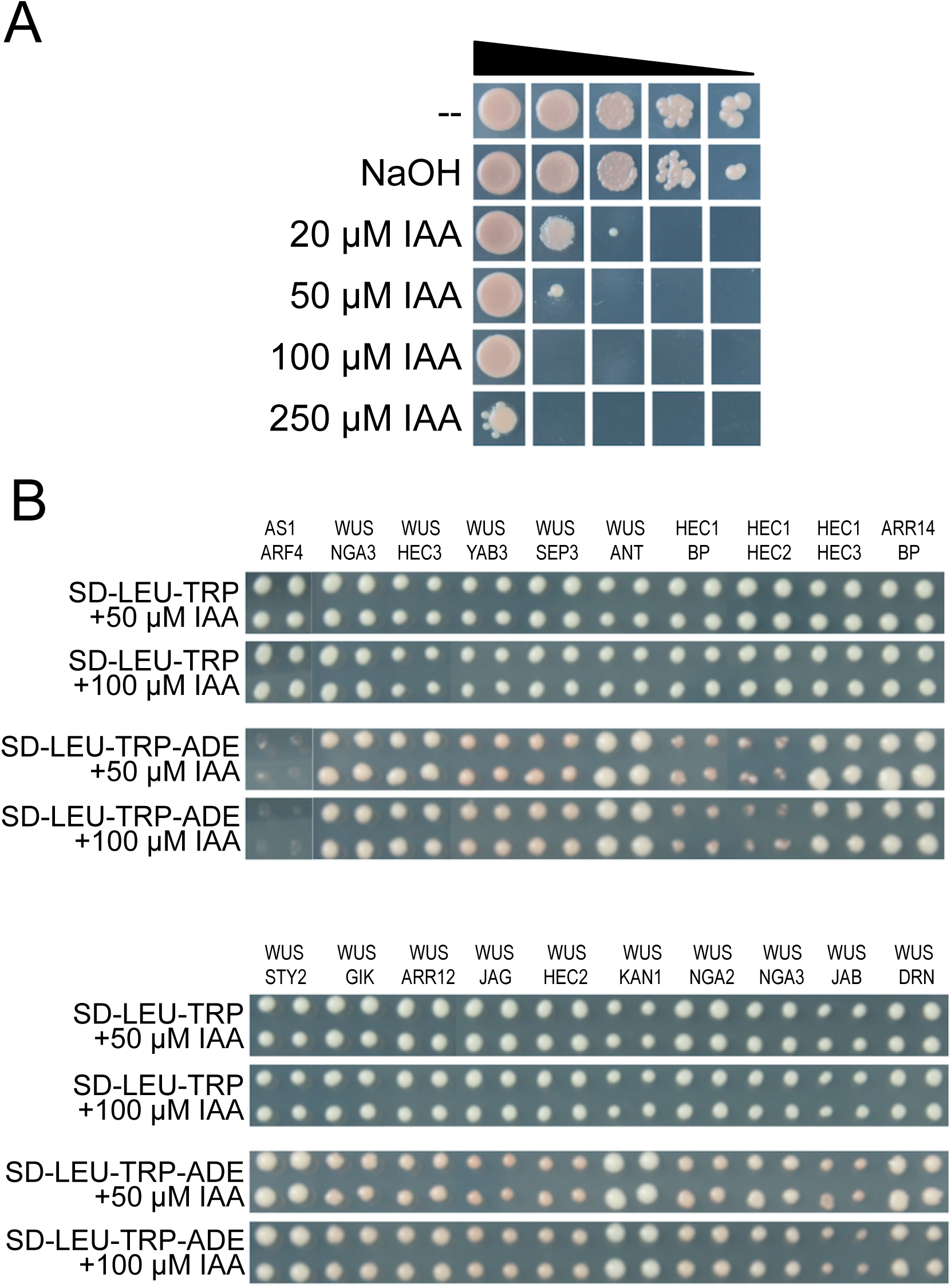
The effect of auxin on protein interactions. Related to Figure 5. A) The addition of IAA to the culture medium does not affect the growth of the PJ69 yeast strain when in high concentrations inoculated. However, a growth effect is observed when dilutions of yeast are inoculated (indicated with the triangle; dilution series 1x, 10x, 100x, 1000x, and 10000x). B) The addition of IAA to the culture medium does not affect the interaction-dependent growth of yeast (high yeast concentration used); some of the tested combinations are presented on medium containing 50 μM and 100 μM IAA.

**Supplemental Figure 5.**
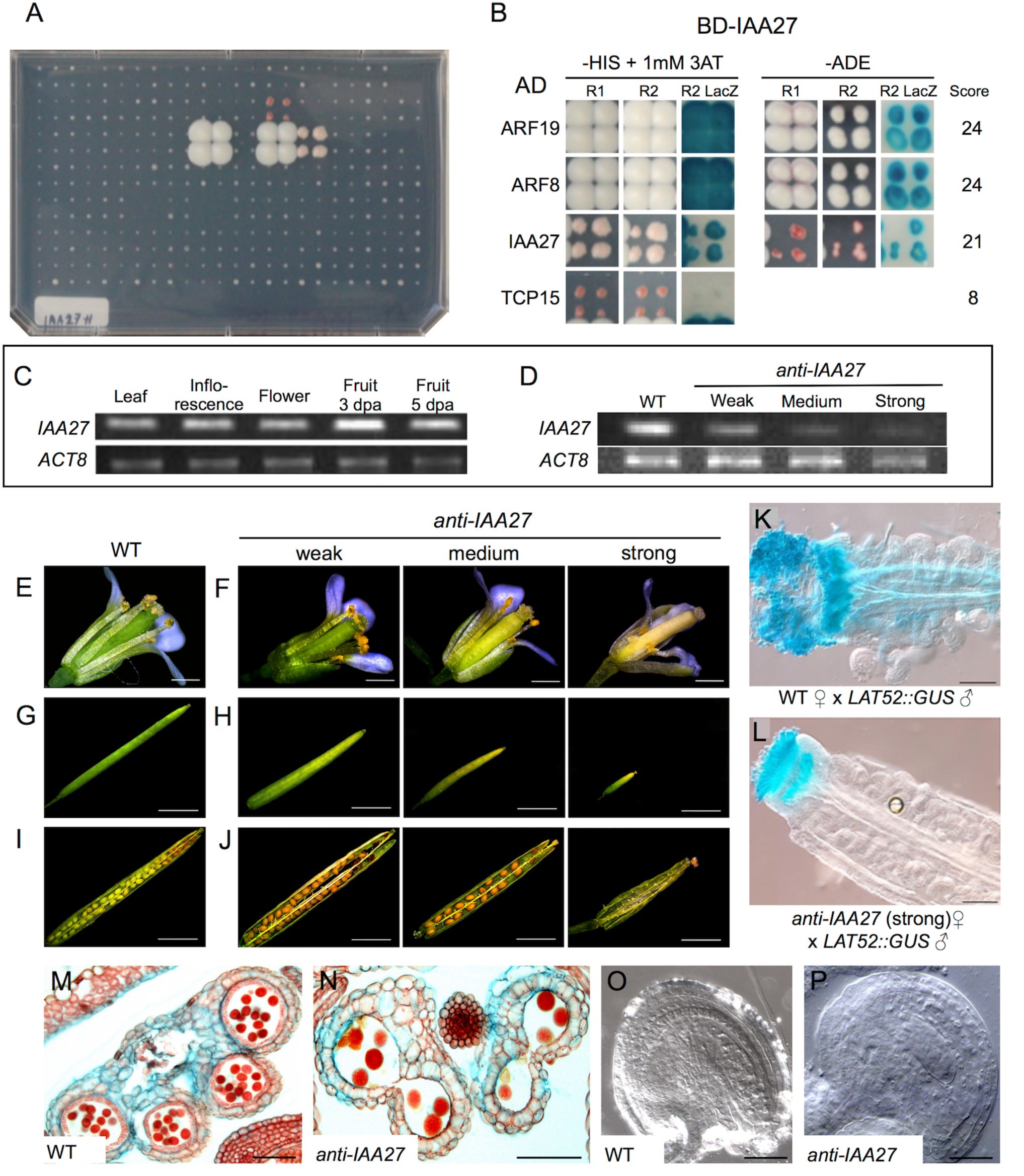
The gene *IAA27* is involved in reproductive development. Related to Figure 6. (A) Overview of the Y2H assay, the plate for BD-IAA27 is shown. (B) Interactions detected for the BD-IAA27 clone based on the 3 interaction markers. (C) Expression of *IAA27* during reproductive development in Arabidopsis. (D) Expression of *IAA27* is reduced at different levels in independent *anti-IAA27* lines. The lines were classified according to the severity of affected fruit size (strong phenotype if the fruit reached up to 35% of its normal size; medium if the fruit reached between 35 and 75% of its normal size, and weak if the fruit reached between 76-90% of its normal size). Flower (E, F) and fruit (G-H) phenotypes of wildtype (WT) plants (E,G,I) and *anti-IAA27* lines (F,H,J). Pollen tube growth visualization in WT (K) and *anti-IAA27* plants (L) after pollination with *LAT52::GUS* pollen. Transverse section of WT (M) and *anti-IAA27* (N) anthers showing that *anti-IAA27* anthers produced fewer pollen grains (pollen degeneration observed) compared with WT anthers. WT ovules (O) compared with *anti-IAA27* (P) ovules showing morphology defects in *anti-IAA27* ovules (affected embryo sac development).

**Supplemental Figure 6.**
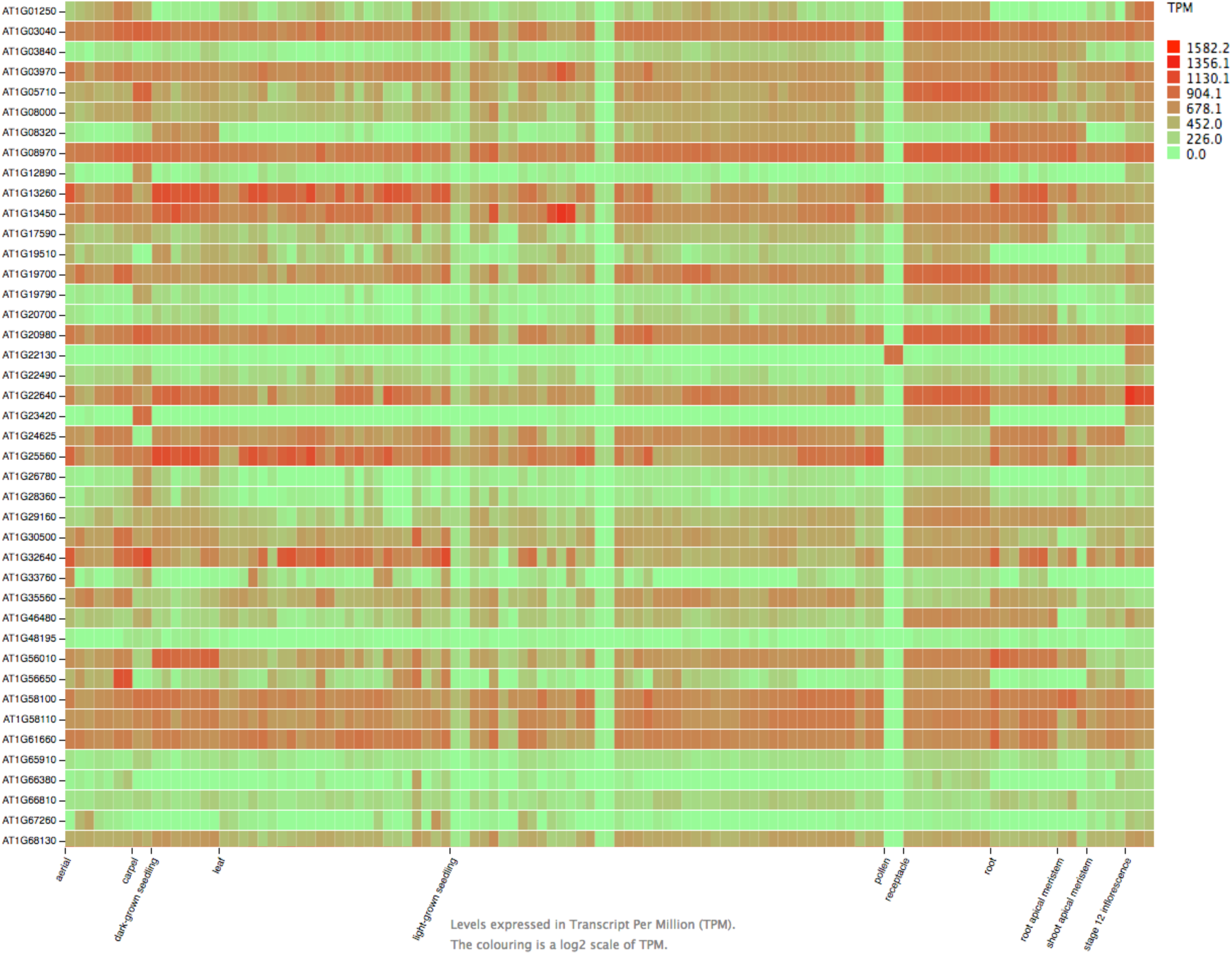
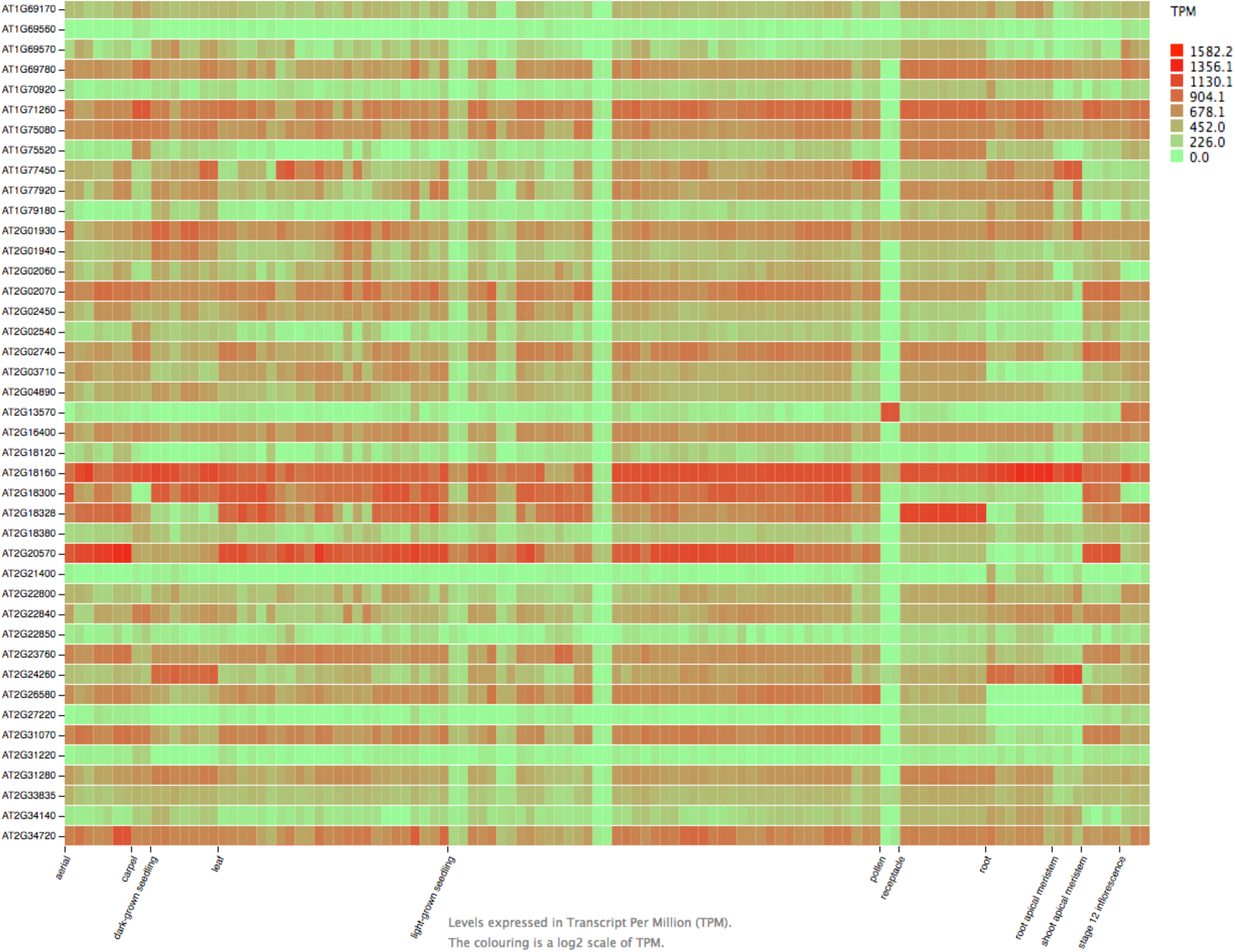
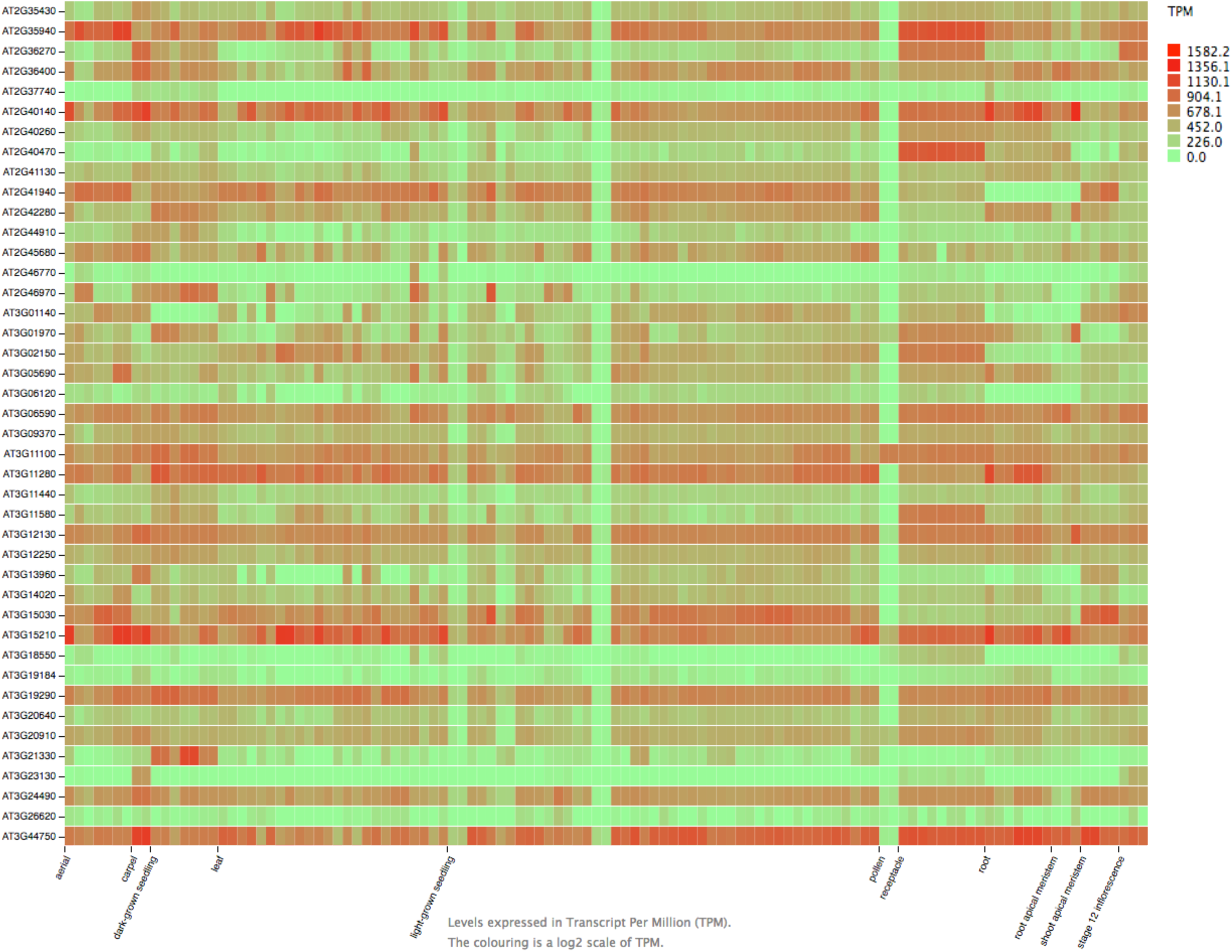
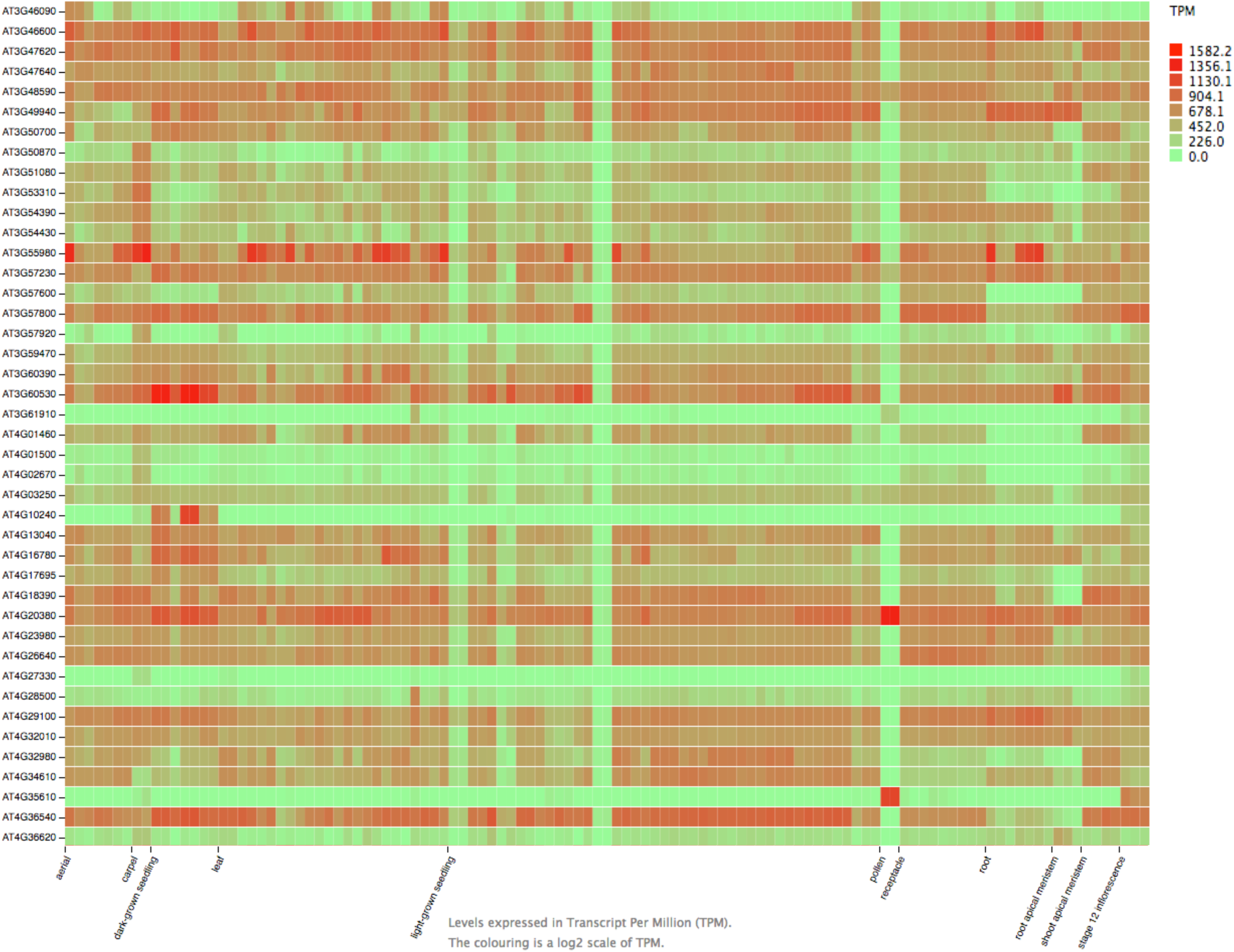
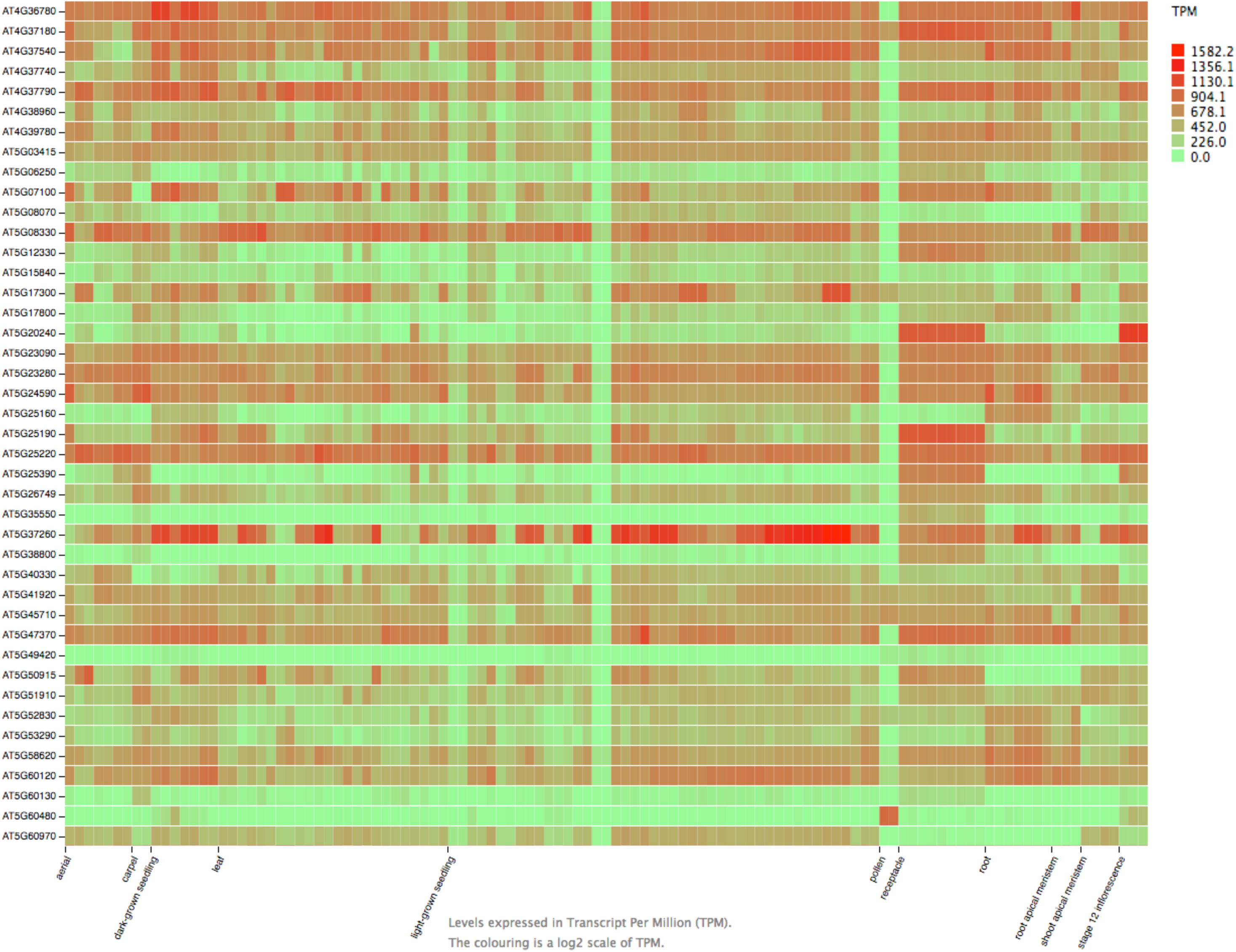
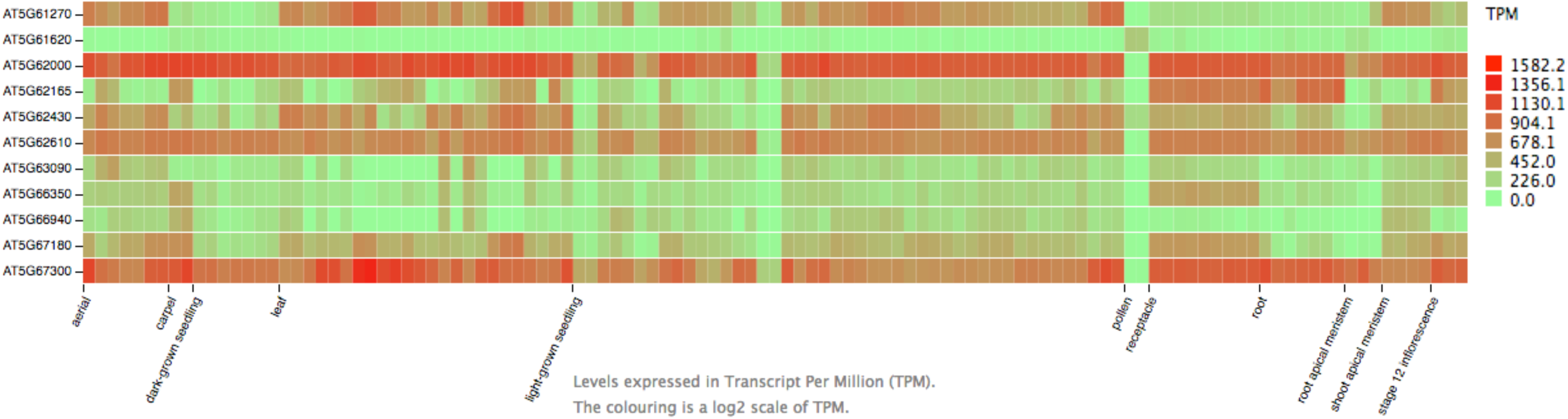
Expression data for the 221 TF candidates of the extended PPI network. Related to Figure 6.

**Supplemental Figure 7. Cytoscape data file for the extended network. Related to Figure 6.**

**Supplemental References.**

